# “Transcriptional and isoform-level regulation of lipid-candidate genes in preeclamptic placentas”

**DOI:** 10.64898/2026.08.17.745256

**Authors:** Kriesha S Eyer, Melanie Lemaire, Xinyu Fan, Samantha L Wilson

## Abstract

Preeclampsia (PE) is a hypertensive pregnancy-specific disorder and a leading cause of maternal and fetal mortality. A common feature of PE placentas and maternal plasma is dyslipidemia, or abnormal lipid levels, which can increase oxidative stress and endothelial dysfunction. However, the precise transcriptional, post-transcriptional, and epigenetic mechanisms underlying these abnormalities remain poorly characterized. Identifying such changes may clarify disease mechanisms and identify lipid-related PE biomarkers. We conducted a large-scale meta-analysis integrating public placental datasets from NCBI GEO, comprising four DNA methylation (DNAm) datasets (n = 172), three RNA-sequencing datasets (n = 92), and an independent RNA microarray validation cohort (n =146). We evaluated differential DNAm (limma), gene expression (DESeq2), transcript-level shifts (Swish), and alternative splicing (rMATS) in PE versus control placentas, with all analyses stratified by fetal sex via an interaction term model. We also performed placental cell-type deconvolution to quantify PE-associated cell-type proportion changes. Our results demonstrated that lipid-related regulation changes in PE placentas occur primarily at the gene and transcript level, with DNAm showing no changes. We also identified significant isoform switching in PE that were undetected by differential gene expression analysis, and primarily driven by alternative transcription initiation and termination sites rather than alternative splicing. A subset of these isoform switches mapped to pathways dysregulated in PE and were predicted to cause functional protein changes. An interaction term model identified several sex-specific differentially expressed genes (DEGs) in PE, including a subset of male-specific downregulated genes involved in oxidative metabolism. However, many of the remaining sex-specific DEGs across both sexes were previously uncharacterized in the literature. These findings suggest that transcriptional and isoform-level regulation play a role in PE-associated dyslipidemia, with certain regulatory pathways displaying fetal sex-specific patterns.

**Highlights:**

- Preeclampsia-associated dyslipidemia manifests at the gene and transcript level
- Reciprocal isoform switches were missed by standard gene-level analyses
- Alternative transcript initiation and termination drove isoform switching
- Sex-interaction modeling identified sex-specific transcriptional shifts in PE

## 1. Introduction

Preeclampsia (PE) is a pregnancy-specific disorder marked by the onset of hypertension and proteinuria after the 20th week of gestation [1]. Affecting 2-8% of pregnancies, PE causes more than 76,000 maternal deaths and 500,000 fetal deaths annually [2–4]. Despite this impact, PE pathogenesis is poorly understood largely due to its heterogeneity. Current literature suggests a multifactorial mechanism involving genetic and environmental factors, maladaptive immune responses, abnormal placentation, reduced cardiovascular adaptation, and endothelial dysfunction [5]. PE also exhibits significant heterogeneity in the timing of onset, affected organ systems, symptom severity, and metabolic changes, making the disorder difficult to predict and treat [6]. Among these metabolic features of PE is dyslipidemia— characterized by the elevation of total cholesterol, triglycerides, and LDL-cholesterol levels rising above the 95th percentile, alongside HDL-cholesterol falling below the 5th percentile for gestational age [6–9].

Lipids are organic molecules that form cell membrane components, and engage in cell signaling, energy storage, heat insulation, and membrane trafficking [10–12]. During healthy pregnancies, the placenta and fetus uptake maternal lipids to meet metabolic and growth demands, produce hormones, and form cell membrane components [13]. However, multiple studies report dyslipidemia in the placenta and maternal plasma of PE pregnancies compared to healthy controls [14,15], characterized by higher levels of total cholesterol, saturated fatty acids, and triglycerides, among other lipid species [16–19]. However, the specific lipid metabolites affected, and the magnitude of this dysregulation vary depending on the timing of disease onset (early-onset vs. late-onset) [20], gestational age at delivery (pre-term vs. term PE) [21], and BMI [22,23]. Metabolic conditions associated with dyslipidemia, such as chronic hypertension, diabetes, and obesity, also act as major PE risk factors [24]. Although, previous studies suggest that the accumulation of triglycerides and fatty acids are correlated with high oxidative stress and endothelial dysfunction in the uterine spiral arteries, seen in PE, the source of this differential lipid regulation and its downstream impact remain unclear [25,26].

Characterizing these lipid-based regulatory changes in PE could reveal mechanisms of disease progression and etiology, identify early lipid-related biomarkers, and inform treatment strategies for lipid- induced PE symptomology. This is especially promising because prediction models using maternal lipid concentrations achieve high accuracy and improve the performance of existing protein biomarkers [27–29]. Abnormal placental lipid metabolism, as seen in PE, has also been implicated in long-term maternal and offspring health conditions like cardiometabolic disease and cognitive impairment [13,30], further highlighting the importance of characterizing the lipid-related changes. However, the relationship between dyslipidemia and PE has sparsely been studied from genetic and epigenetic perspectives which may help identify the regulatory mechanisms influencing PE risk downstream cardiometabolic disease. To address this gap, we examined lipid-related gene patterns through an RNA expression, alternative splicing, and DNA methylation (DNAm) lens.

RNA expression analysis provides information about the types and amounts of RNA species in tissues, revealing how protein output or cellular function may change between diseased and healthy states [31]. Previous studies show that PE placentas exhibit large expression shifts in genes governing inflammation, hypoxia, oxidative stress, angiogenesis, and lipid metabolism [32].

Beyond total expression levels, one source of genetic change that is of increasing interest in PE is differential isoform usage, and alternative splicing [33,34]. Alternative splicing involves combining different exons from a single mRNA transcript to produce isoforms and proteins with novel functions [35]. For example, transcripts involved in lipid uptake, synthesis, and oxidation may undergo alternative splicing to alter protein activity or localization [36,37]. In some diseases, these newly generated isoforms may be less functional, driving pathology independent of total gene expression changes by shifting the ratio of functional to non-functional isoforms [38,39]. It is, therefore, possible that isoform shifts in the PE placenta may be a source of the observed dyslipidemia, if changes in total transcript abundance are not detected.

Both RNA expression and alternative splicing can be regulated by epigenetic modifications, including DNAm—the addition of a methyl group to the 5’ position of a cytosine nucleotide next to a guanosine (CpG sites) in the DNA sequence [40]. DNAm typically represses gene expression by interfering with transcription factor binding [40]. In alternatively spliced exons, DNAm also regulates exon inclusion levels, dictating the predominant isoforms of a given gene [41]. Therefore, differential epigenetic signatures in PE pregnancies may alter both gene expression and splicing patterns to drive metabolic changes [42].

The objective of this study was to conduct a meta-analysis of publicly available placental RNA-sequencing and DNAm data, to assess altered gene expression, alternative splicing, differential isoform usage, and DNAm patterns in PE. We first conducted targeted analyses on a candidate list of lipid-related genes to increase statistical power, as genome-wide analyses may miss subtle expression changes due to stringent multiple test corrections. We then repeated the analyses genome-wide to identify novel altered pathways beyond lipid metabolism. Given the known sex differences in PE timing of onset and prevalence [43,44], we completed both combined-sex and sex-stratified analyses via an interaction term model. This allowed us to explore whether genetic and epigenetic changes differ between female and male fetuses. Our workflow utilized linear modeling and functional pathway enrichment using the Gene Ontology (GO) database. We hypothesized that RNA expression, isoform usage, and DNAm changes in lipid-related genes are prominent and sex-specific in PE.

## 2. Material and Methods

### 2.1 Lipid-Candidate Gene Identification

We retrieved lipid-related genes from the LIPID MAPS® Gene/Proteome Database (LMPD) [45], and DBLiPro databases [46], both of which contain records on proteins involved in human lipid using annotations from UniProt, KEGG, and Gene Ontology. We supplemented this base list with lipid-related genes from the Genetic Association Database, which links specific genes to human diseases [47], and the Reactome Pathways 2024 database [48–50], which contains genes involved in known human molecular pathways and reactions. To filter these databases for lipid-related targets, we applied a search term string adapted from the Comprehensive Lipid Classification Framework [51]. Search terms are available in our GitHub repository [52]. To further identify lipid pathology related candidate genes, we manually searched peer-reviewed studies focused on lipid and metabolic disorders, including preeclampsia [53–62]. Merging these lists and removing duplicates yielded 5,510 genes. We matched these gene names against ENSEMBL IDs from GENCODE v48 to enable subsequent filtering of the RNA-seq data [63]. We excluded genes with no corresponding ENSEMBL IDs, leaving 5,321 unique lipid-candidate genes (Supplementary Table 1).

### 2.2 GEO Screens/Data Download for RNA-Seq Data

This study adheres to the guidelines outlined in the Preferred Reporting Items for Systematic Reviews and Meta-Analyses (PRISMA) 2020 statement (Figure 1a) [64]. We searched the NCBI Gene Expression Omnibus (GEO) database using the keywords “preeclampsia” and “RNA”. Our inclusion criteria required paired-end, Illumina bulk RNA-seq data from singleton, full-term chorionic villi samples that included both preeclamptic and healthy controls. Paired-end read lengths ranged between 124 bp and 150 bp depending on the study group. We downloaded raw fastq files for GSE143953 [65], GSE148241 [66,67], GSE186257 [68], GSE234729 [69], GSE255126 [70], and GSE279757 [71], using the Sequence Read Archive (SRA) Toolkit v3.2.1 “fasterq-dump --split-files” command, which yielded 228 starting samples [72]. We retrieved the corresponding clinical metadata using the GEOquery R package v2.72.0 [73]. All data processing and analysis were performed using R (v.4.4.2) [74].

**Figure 1:**
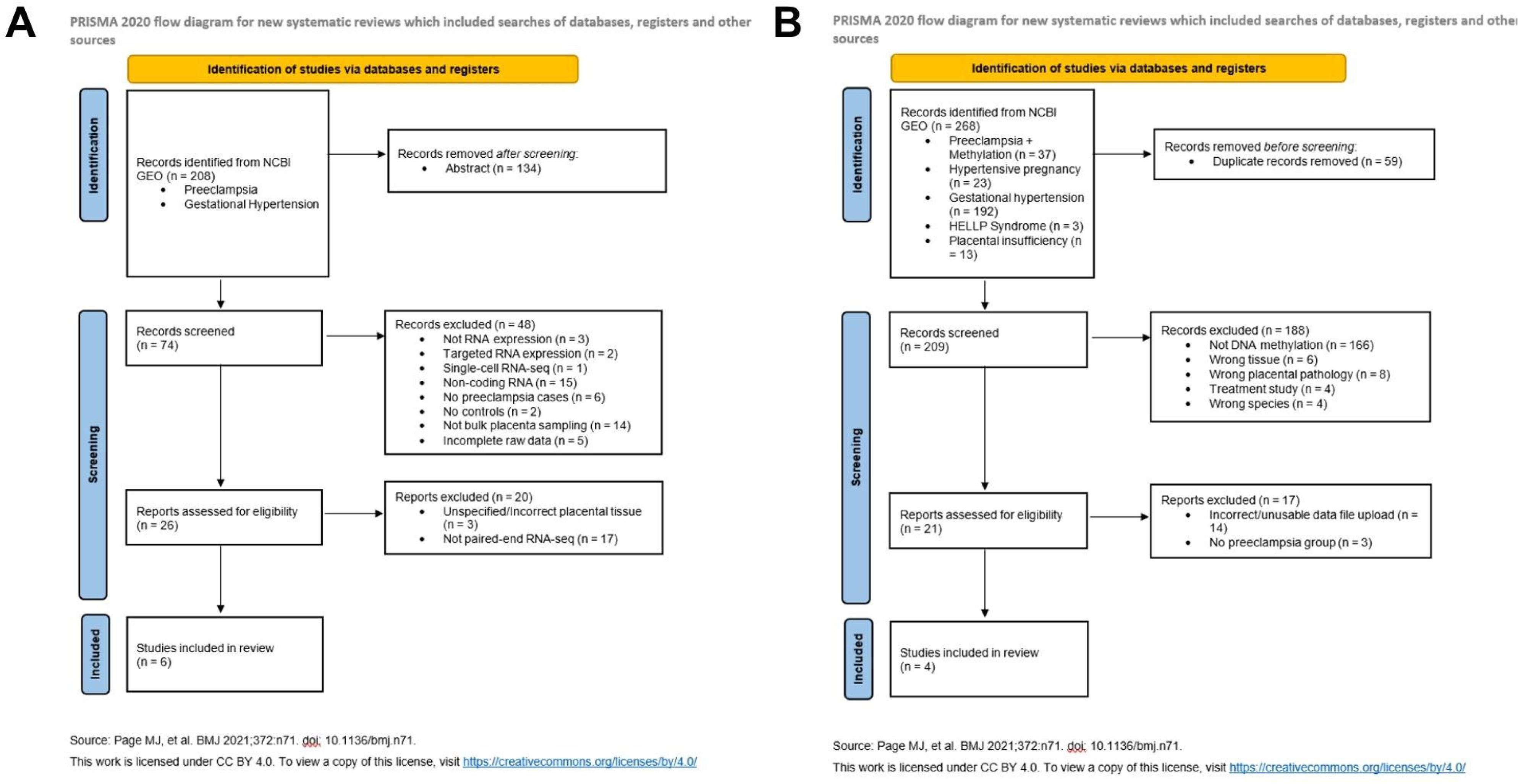
Flowchart of dataset selection of placental RNA-sequencing datasets (A) and placental DNAm Illumina Methylation 450K array datasets (B) from GEO based on inclusion and exclusion criteria, assembled using the PRISMA 2020 guidelines [64].

### 2.3 RNA-Seq Quality Control and Alignment

We assessed initial read quality using FastQC v0.12.0 [75], and FastQ Screen v0.14.0 [76]. We trimmed low-quality bases and removed sequencing adapters using fastp v0.23.4 [77]. To quantify library duplication rates, we utilized fastp v0.23.4 [77], picard’s EstimateLibraryComplexity v3.4.0 [78], and the dupRadar R package v1.34.0 [79]. We excluded samples exhibiting high duplication, defined as either a dupRadar y-intercept greater than 50% or an estimated library complexity below 10 million unique reads (n = 24), leaving 204 samples. We identified duplicate reads using Picard’s markDuplicates v3.4.0 [78] and aligned the reads to the reference genome using STAR v2.7.11b [80], discounting duplicate reads before quantification. We excluded one sample (SRR16505233 from GSE186257) from downstream analyses due to sequencing quality issues, with only 1.7% of reads mapped to the reference genome, leaving 203 samples before batch effect analysis. Supplementary Figure 1 outlines a flowchart for all samples removed. We predicted fetal sex using the SexInference R package v1.2.0 from SexChrLab [81]. All algorithmic predictions matched the reported clinical sex.

### 2.4 RNA-Seq Batch Effect Analysis

We performed principal component analysis (PCA) using the ggplot2 R package v3.5.1[82] to assess batch effects across the normalized datasets (GSE143953, GSE148241, GSE186257, GSE234729). GSE234729 samples formed two distinct, isolated clusters separated from the other cohorts which standard batch-correction methods could not resolve (Supplementary Figure 1). To eliminate this technical variation, we excluded all 111 samples from GSE234729, leaving 92 samples for downstream analysis. Supplementary Figure 2 visualizes the combined RNA-seq datasets before and after normalization to confirm successful batch effect removal. Table 1 displays the final sample demographics.

**Table 1:**
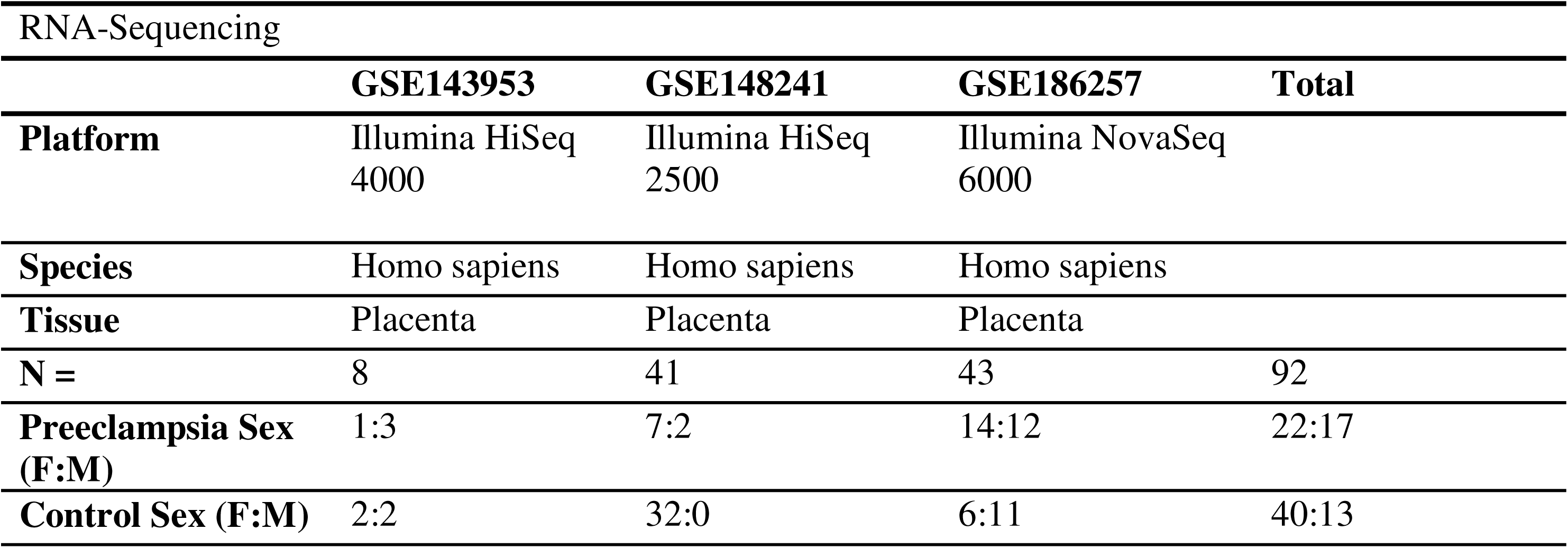
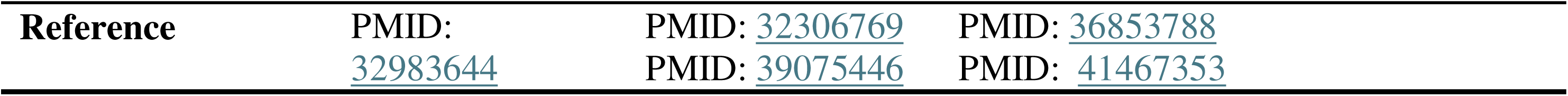
Sample demographics for 3 RNA-sequencing studies obtained from GEO used in differential gene expression meta-analysis.

| RNA-Sequencing |  |  |  |  |
| --- | --- | --- | --- | --- |
|  | GSE143953 | GSE148241 | GSE186257 | Total |
| **Platform** | Illumina HiSeq 4000 | Illumina HiSeq 2500 | Illumina NovaSeq 6000 |  |
| **Species** | Homo sapiens | Homo sapiens | Homo sapiens |  |
| **Tissue** | Placenta | Placenta | Placenta |  |
| **N =** | 8 | 41 | 43 | 92 |
| **Preeclampsia Sex (F:M)** | 1:3 | 7:2 | 14:12 | 22:17 |
| **Control Sex (F:M)** | 2:2 | 32:0 | 6:11 | 40:13 |

|  |  |  |  |
| --- | --- | --- | --- |
| **Reference** | PMID: 32983644 | PMID: 32306769<br>PMID: 39075446 | PMID: 36853788<br>PMID: 41467353 |

### 2.5 RNA-Seq Differential Expression Analysis

We filtered the RNA read count matrix to include only the 5,321 lipid-candidate ENSEMBL IDs identified in Section 2.1. The DESeq2 R package v1.52.0 modeled expression differences between the PE and control groups using negative binomial generalized linear models, adjusting for study group and fetal sex [83]. We analyzed autosomes first in the combined population, followed by sex-stratified subsets. Next, we incorporated an interaction term (fetal_sex*pathology_group) into the design formula to compare its results to the sex-stratified models, and identify whether fetal sex significantly alters the effect of PE on gene expression [83].

We modeled X-chromosome expression separately for each sex. We excluded the Y-chromosome from subsequent analyses as none of the identified lipid candidate genes mapped to this chromosome. To correct for multiple testing, we applied the Benjamini-Hochberg false discovery rate (FDR) method, with significance defined as an FDR < 0.05 [84]. We subsequently repeated this methodology genome-wide.

### 2.6 Gene Ontology and Pathway Enrichment Analysis

We performed precision-recall scoring using the ErmineJ R package v.3.1.2 to conduct pathway enrichment across the differentially expressed genes (DEGs) [85]. We utilized Gene Ontology annotations from the Gemma database, to reveal possible overrepresented functional pathways among our DEGs [86,87]. Significant pathways required an FDR < 0.05 and a multifunctionality score < 0.5. Because a high multifunctionality score reflects genes annotated to many functional pathways, this correction ensures that the identified pathways are driven by genes specific to a distinct biological process [85].

### 2.7 RNA-Based Cell-Type Deconvolution

To estimate and compare placental cell proportions between the pathology groups, we performed RNA reference-based cell-type deconvolution using the MuSiC R package v1.0.0 [88,89]. We utilized the single-cell RNA-sequencing dataset GSE182381 as the reference matrix to define baseline placental cell-type signatures [90]. The algorithm calculated proportions of cytotrophoblasts, endothelial cells, extravillous trophoblasts, fibroblasts, GZMB/GZMK natural killer cells, hofbauer cells, mesenchymal stem cells, nucleated red blood cells, proliferative cytotrophoblasts, and syncytiotrophoblasts across the combined dataset.

### 2.8 Affymetrix Differential RNA Expression Validation Cohort

To validate our findings on an independent platform, we analyzed placental RNA data from a Affymetrix Human Gene 1.0 ST Array. We downloaded raw .CEL files from the GSE75010 dataset (80 PE, 77 non-PE) [6], and processed them using the Biobase v2.70.0 [91], and oligo v1.74.0 [92] R packages to normalize the data, filter out lowly expressed genes, and remove outliers, following the “Affymetrix end-to-end workflow” [93]. The arrayQualityMetrics v3.66.0 R package [94] identified eleven samples with outlier signal intensity distributions indicating array quality issues. Excluding these samples left 74 PE and 72 control samples for analysis. We annotated transcript clusters using the AnnotationDBI R package v1.72.0 [95]. Sample demographics are displayed in Table 2.

**Table 2:**
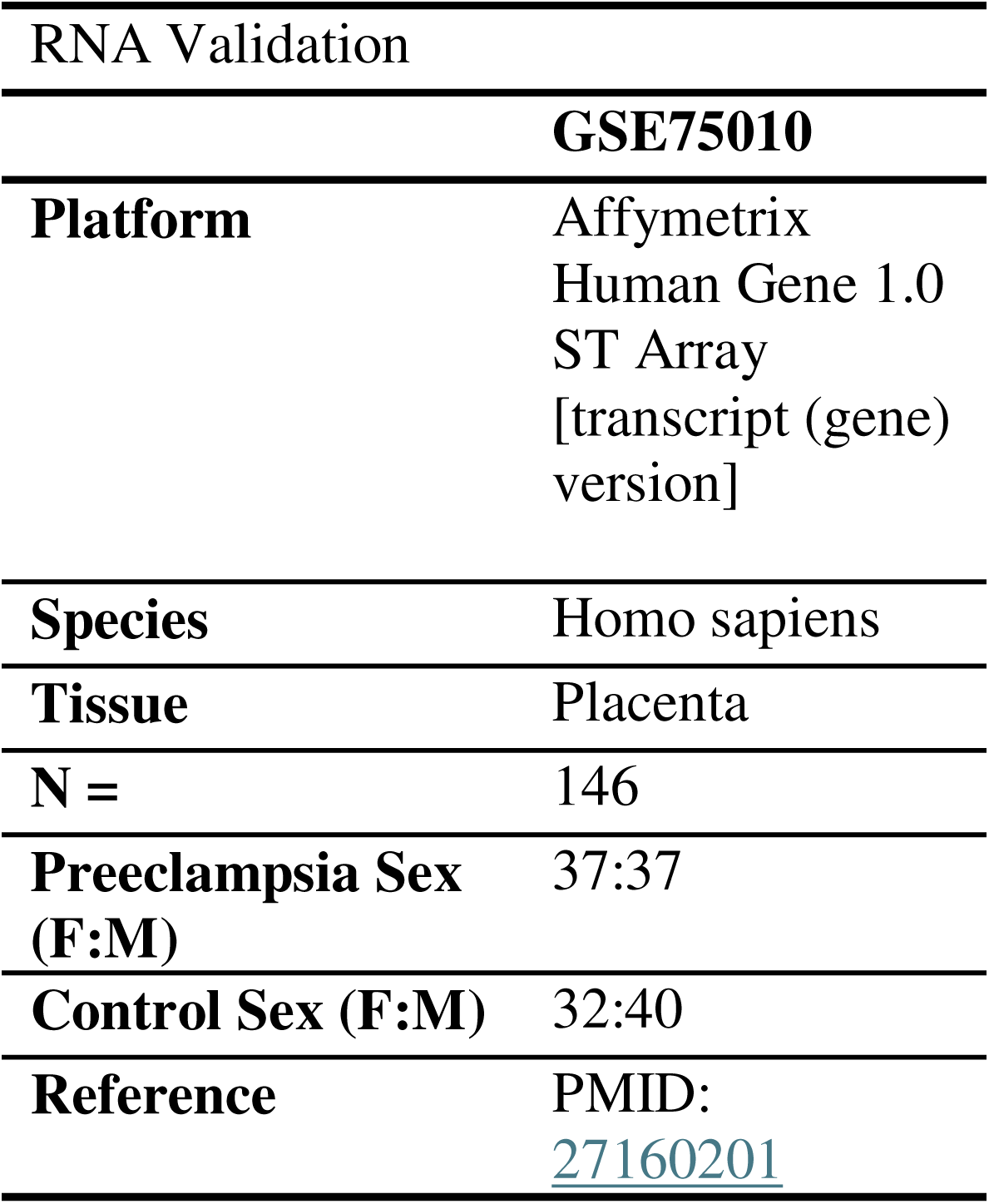
Sample demographics for Affymetrix Validation cohort obtained from GEO.

We subset this dataset to contain only the significant DEGs identified in our primary RNA-seq analyses from the combined-sex population and the fetal-sex interaction term model (Section 2.5). We then performed differential expression modeling using the limma R package v3.60.6, adjusting for gestational age and fetal sex [96]. We analyzed the combined population first, followed by sex-stratified subsets. Significance was set at an FDR < 0.05. We determined the percent validation by dividing the number of successfully validated genes in the Affymetrix dataset by the total number of genes common to both RNA-seq and Affymetrix datasets.

### 2.9 Alternative Splicing and Isoform Switching Analysis

To capture transcript-level variations missed by gene-level DESeq2 modeling, we mapped differential isoform expression using the Swish R package v1.6.0 [97]. We calculated transcript abundances by mapping raw fastq files to the GENCODE v48 reference genome using Salmon v1.10.3 with 30 bootstrap iterations [63,98]. Swish modeled transcript variation across the combined and sex-stratified cohorts. Because software limitations prevented us from combining batch correction and interaction modeling, we analyzed sex-specific differential isoform expression using standard sex-stratified cohorts. Because the male GSE148241 cohort contained two PE samples and zero controls, we removed these two PE samples prior to analysis. We also removed three female samples from GSE143953 because that cohort contained only two controls and one PE sample, leaving the total sample set for differential isoform analysis to be 87 samples (51 control: 38 female, 13 male; 36 PE: 21 female, 15 male).

To identify the alternative splicing events driving the transcript-level variations within a single gene, we ran the rMATS-turbo package v4.0.1 [99] on the initial STAR-mapped BAM files. We restricted rMATS analysis to the whole-genome data of the combined-sex population since the sex-stratified analyses lacked sufficient statistical power.

We ran the IsoformSwitchAnalyzeR v2.12.0 [100–102] and DEXSeq v1.58.0 [96,100,103] R packages on the Salmon output files to identify preferred isoform switches between PE and control groups. We evaluated these switches within the lipid-candidate genes and genome-wide across the combined and sex-stratified populations. We evaluated the downstream functional consequences of these structural switches including protein coding potential using CPC2 [104], protein domain changes using PFam [105], signal peptide adjustments using SignalP v6.0 [106], and disordered region changes using IUPred2A v3.0 [107].

### 2.10 GEO Screens/Data Download of DNAm Data

We again followed the guidelines of the PRISMA 2020 statement (Figure 1b) [64]. We identified placental DNAm datasets by searching the NCBI GEO database using the keywords "preeclampsia," "methylation," "hypertensive pregnancy," “HELLP syndrome,” and "placental insufficiency." This search yielded 4 datasets that met our criteria: GSE100197 [108], GSE98224 [108,109], GSE75196 [110], and GSE125605 [111]. All included studies utilized the Illumina Infinium HumanMethylation450k BeadChip array on full-term, singleton chorionic villi samples, and contained both PE and control samples.

We downloaded raw intensity (IDAT) files and retrieved corresponding metadata using the GEOquery R package v2.72.0 [73]. We used the minfi R package v1.50.0 to predict fetal sex as a quality control measure and to provide missing fetal sex metadata for the GSE75196 dataset [112]. All reported sexes matched minfi predictions. From the GSE100197 dataset, we excluded samples of IUGR pathology (n = 11), pre-term PE pregnancies (n = 24), and technical replicates (n = 8) since they were available only for this study. We also excluded one sample from GSE125605 (n = 1) due to missing gestational age data. Final sample demographics are displayed in Table 3.

**Table 3:** Sample demographics for 4 DNA methylation studies obtained from GEO used in differential DNA methylation meta-analysis.

|  |  |  |  |  |  |
| --- | --- | --- | --- | --- | --- |
| DNA Methylation |  |  |  |  |  |
|  | **GSE75196** | **GSE98224** | **GSE100197** | **GSE125605** | **Total** |
| **Platform** | Illumina<br>Human 450K<br>Methylation<br>Array | Illumina<br>Human 450K<br>Methylation<br>Array | Illumina<br>Human 450K<br>Methylation<br>Array | Illumina<br>Human 450K<br>Methylation<br>Array |  |
| **Species** | Homo sapiens | Homo sapiens | Homo sapiens | Homo sapiens |  |
| **Tissue** | Placenta | Placenta | Placenta | Placenta |  |
| **N =** | 24 | 48 | 59 | 41 | 172 |
| **Preeclampsia Sex (F:M)** | 4:4 | 16:14 | 16:24 | 12:10 | 48:52 |
| **Control Sex (F:M)** | 9:7 | 6:12 | 10:9 | 10:9 | 35:37 |
| **Reference** | PMID: 26968548 | PMID: 29507646<br>PMID: 29092053 | PMID: 29092053 | PMID: 31091979 |  |

### 2.11 DNA Methylation Data Normalization and Probe Filtering

We performed functional normalization on the combined IDAT using minfi v1.50.0, utilizing control probes to minimize background noise and technical batch variations between independent experiments [113]. We constructed pre- and post-normalization PCA plots via ggplot2 v3.5.1 to track alignment quality, using missMDA v1.20 to estimate 1717 missing methylation values in the raw data (Supplementary Figure 4) [82,114]. The normalized output removed missing DNA methylation values.

Following normalization, probe filtering removed probes with missing beta values or detection p-values > 0.01 in more than 5% of samples. We annotated sex chromosomes using the IlluminaHumanMethylationEPICanno.ilm10b4.hg19 R package, zeroing positive Y-chromosome signals in female samples to prevent artifacts [115]. Finally, we excluded cross-hybridizing probes, SNP-associated probes, and non-variable placental probes to prevent false-positive signals [116].

### 2.12 Differential DNAm Analysis of Candidate Lipid-Related Genes and Genome-Wide Loci in PE

We mapped Illumina CpG probe IDs to gene names by aligning CpG genomic position to the UCSC hg38 reference genome, then filtered this output for our candidate lipid-candidate gene. We also annotated the closest transcription start site to each CpG site using an annotation file from Price et al. [117]. We analyzed differential DNAm between the PE and control groups using the limma R package v3.60.6 [96] 4across combined autosomal, sex-stratified autosomal, and separate X-chromosome datasets.

We performed linear modeling using the “lmFit” and “eBayes” function in limma, adjusting for gestational age and fetal sex, performing the analysis both with ((∼pathology_group*Fetal_Sex + GSE_number + gestational_age) and without ((∼pathology_group + Fetal_Sex + GSE_number + gestational_age) an interaction term [118]. We corrected for multiple testing using the Benjamini-Hochberg FDR method [84]. Significant differentially methylated CpGs required an FDR < 0.05 and an absolute delta-beta value greater than 0.05 (|Δβ| > 0.05), where positive values represent an increase in DNA methylation and negative values reflect a decrease in DNA methylation. We generated volcano plots using ggplot2 [82]. We conducted this analysis on the lipid-candidate genes first, followed by a genome-wide assessment. We then applied the ErmineJ precision-recall pathway enrichment workflow described in Section 2.6 to these methylation results [85]. We compiled genes that features differences in DNAm, RNA expression in RNA-seq datasets, and RNA expression in the Affymetrix dataset to identify the genes with the most prominent epigenetic and genetic changes in PE placentas.

### 2.13 Analysis of Differentially Methylated Regions (DMRs) in Candidate Lipid-Related Genes and Genome-Wide Loci in PE

We used the DMRcate R package v 3.0.10 [119] to identify clustered regions of DNAm (DMRs) across both the lipid-candidate genes and genome-wide. Because regional epigenetic shifts are more likely to interfere with transcriptional machinery than individual CpG changes, DMRs have a higher probability of altering the phenotype of gene expression or protein concentration [120]. The analysis utilized a 1000- nucleoside bandwidth and a scaling factor of 2. We modeled regional adjustments on combined autosomes, sex-stratified autosomes, and individual sex chromosomes separately. Significant regional switches required a harmonic mean of the individual CpG p-values (HMFDR) < 0.05 and an absolute average regional methylation shift greater than 0.05 (∣mean Δβ∣>0.05).

### 2.14 DNAm-based Placental Cell Deconvolution for PE and Control Samples

To estimate and compare cell-types proportions in the bulk placental tissue between PE and control samples, we performed cell deconvolution using the planet R package v1.20.0 based on the DNA methylation dataset [121]. Utilizing reference placental DNAm signatures from the EpiDISH R package v2.28.0 [122], the algorithm calculated individual sample proportions for trophoblasts, stromal cells, hofbauer cells, endothelial cells, nucleated red blood cells (nRBCs), and syncytiotrophoblasts [123]. We compared these cell-type proportions across pathology group and fetal sex using a mixed-effects model via the lme4 R package v1.1.35.4, treating study (GSE number) as a random effect [124].

## 3. Results

### 3.1 Autosomal analyses reveal differential expression in lipid-candidate genes and genome-wide in PE placentas

To evaluate the gene expression differences between healthy and PE placentas, we performed differential gene expression analysis on our lipid-candidate gene list. In the unstratified, combined-fetal-sex population, 14 lipid-related genes were significantly downregulated (FDR < 0.05, log2FC < -1) and 62 genes were significantly upregulated genes (FDR < 0.05, log2FC > 1; Figure 2a). Top upregulated and downregulated lipid-candidate genes are noted in Supplementary Table 2.

**Figure 2:**
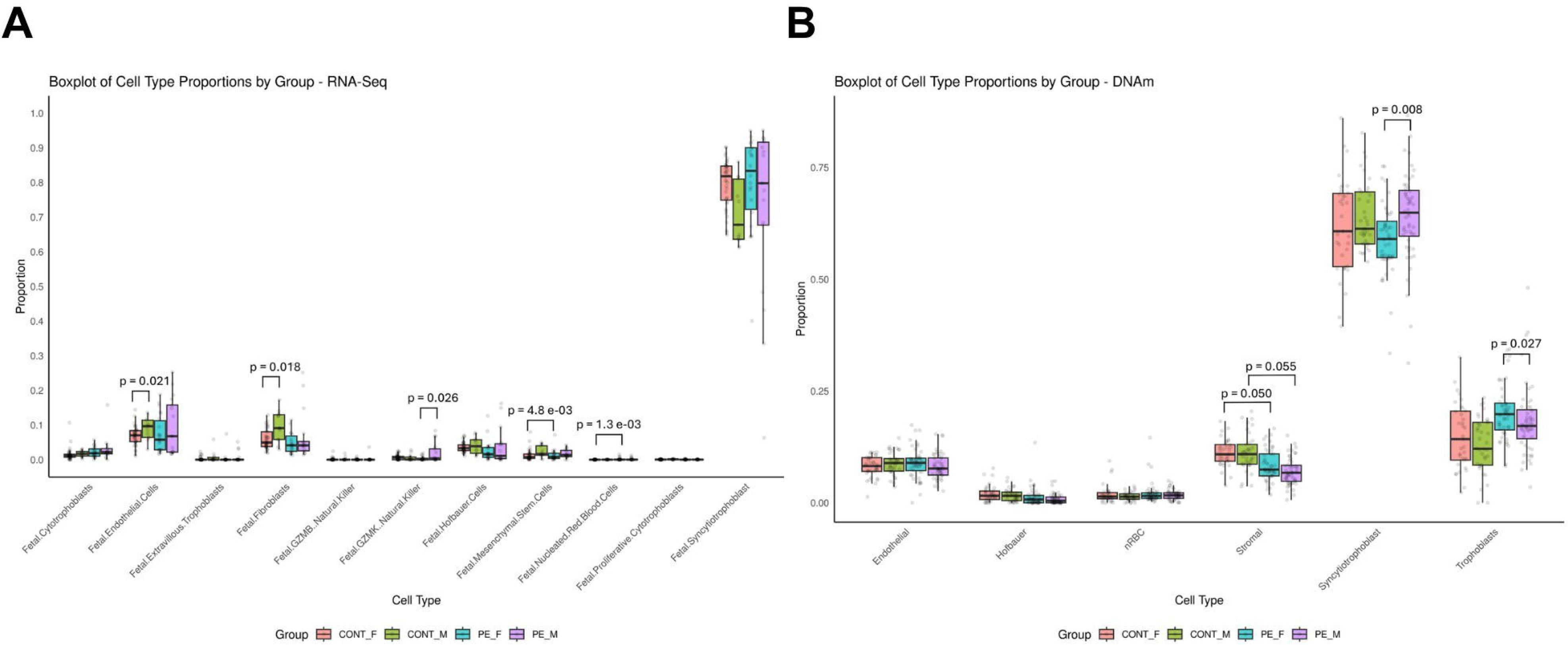
Volcano plot depicting autosomal differential lipid-related gene expression (RNA-Seq) between PE and control placentae in the (A) combined fetal sex population (n = 92; 53 PE, 39 Control), (B) female-stratified population (n = 62; 22 PE, 40 Control), and (C) male-stratified population (n = 30; 17 PE, 13 Control). The x-axis represents the magnitude of signal change, with higher absolute values indicating greater changes. The y-axis shows the FDR, where higher values reflect greater statistical significance. Purple points represent genes with biologically significant decreased expression in PE, while pink points indicate genes with biologically significant increased gene expression in PE. Grey points represent genes with no significant differential gene expression.

We expanded the differential gene expression analysis genome-wide to capture non-lipid related changes. In the combined-fetal-sex autosomal population, 450 genes showed significant upregulation and 148 genes showed significant downregulation in PE placentas (Figure 3a). Full genome-wide differential expression results are provided in Supplementary Table 3.

**Figure 3:**
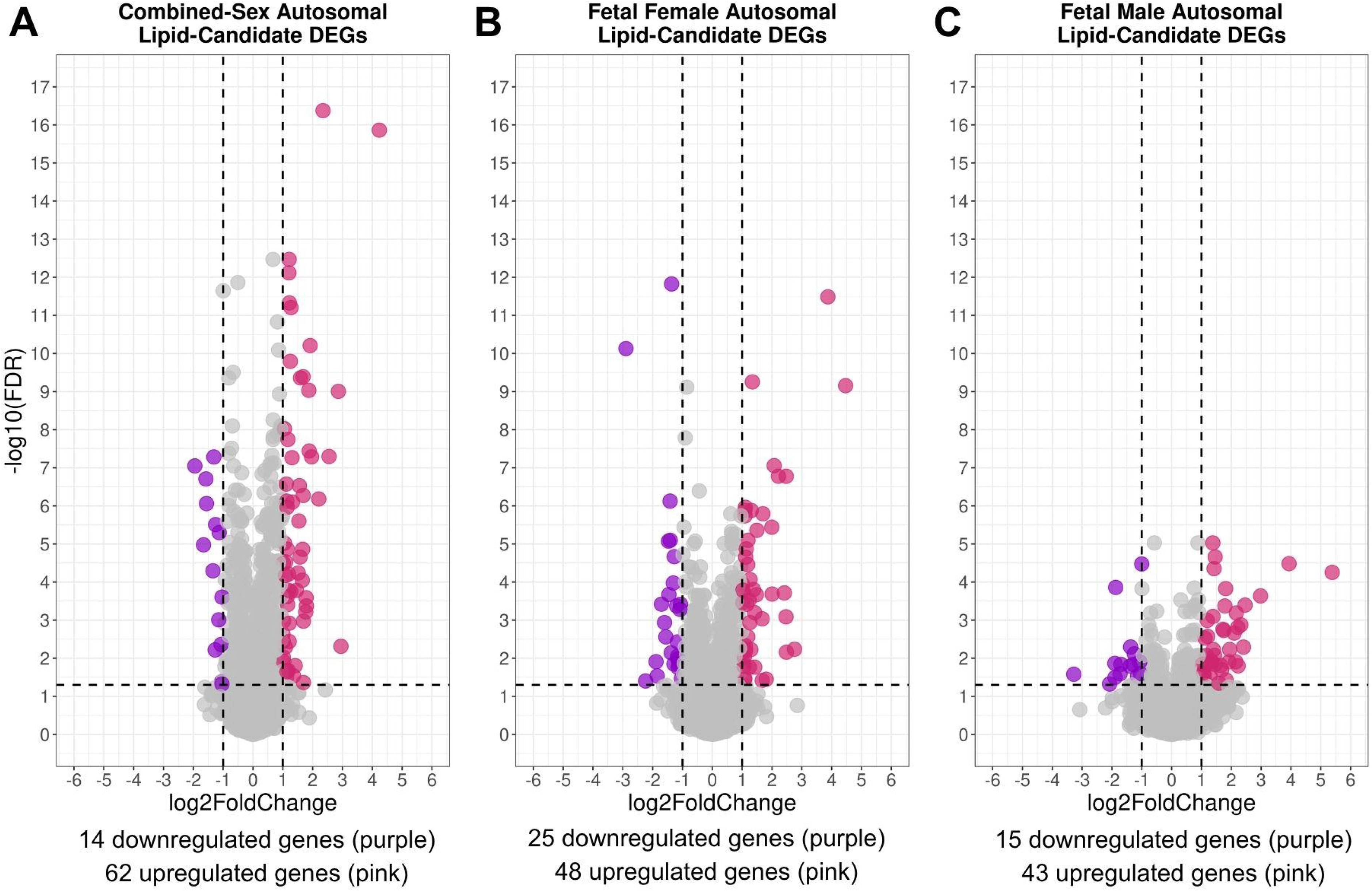
Volcano plot depicting autosomal differential genome-wide gene expression (RNA-Seq) between PE and control placentae in the (A) combined fetal sex population (n = 92; 53 PE, 39 Control), (B) female-stratified population (n = 62; 22 PE, 40 Control), and (C) male-stratified population (n = 30; 17 PE, 13 Control).

### 3.2 Interaction term modeling increases statistical power and reveals sex-specific differentially expressed genes in PE placentas

To identify possible sex-specific differential gene expression patterns in PE, we performed and compared the results of sex-stratified (male-only and female-only) analyses and a fetal-sex-pathology interaction term model. While sex-stratification splits the datasets by sex and lowers statistical power, the interaction term model uses all samples to calculate background variance and prevent false negatives. In the lipid- candidate analysis, sex-stratification revealed 48 significantly upregulated and 25 downregulated genes in fetal female autosomes in PE (Figure 2b), compared to 43 upregulated and 15 downregulated genes in fetal male autosomes (Figure 2c). Thirty-one genes were commonly differentially expressed across both fetal sexes.

Analysis of X-chromosomal lipid-related genes revealed 1 up-regulated and 1 down-regulated gene in fetal females, compared to only 1 up-regulated gene in fetal males in PE (Supplementary Figure 5). The interaction term model yielded similar trends to the sex-stratified approach in the female-specific interaction analyses, with 51 genes showing increased expression and 24 genes showing decreased expression. However, in the fetal male autosomal interaction analysis there were almost double the number of significant DEGs, with 79 upregulated genes and 27 downregulated genes in PE (Supplementary Table 2). Among these DEGs, only 19 were commonly differentially expressed across both fetal sexes. There were no hits in the interaction term model for the X-chromosomal lipid-related genes.

In the whole-genome analysis, sex-stratification revealed 336 upregulated and 193 downregulated genes in fetal females (Figure 3b), and 242 upregulated and 64 downregulated genes in fetal males in PE (Figure 3c), with 91 shared between fetal sex. The X-chromosomal analyses revealed 12 upregulated and 12 downregulated genes in female PE placentas, compared to 7 upregulated and 2 downregulated genes in male PE placentas (Supplementary Figure 6). The interaction term model aligned with the sex-stratified approach in females, with 312 genes showing significant upregulation, and 194 genes showing downregulation (Supplementary Table 3). Similar to the trends in the lipid-candidate analyses, the male whole-genome interaction model identified a higher number of DEGs, with 461 upregulated and 172 downregulated genes (Supplementary Table 3). Among these DEGs, only 124 were commonly differentially expressed across both fetal sexes. There were no hits in the interaction term model for the X-chromosomal genes.

The interaction term model also identified genes with significant expression differences between females and males in PE. In the lipid-candidate analysis, *IGKC* and *LDHB* showed a significant, male-specific decrease in expression in PE placentas (FDR < 0.05, ∣log2FC∣>1; Table 4). In the whole-genome analysis, the interaction term model identified 26 DEGs exhibiting sex-specific regulation in PE. Among these, *IGKC*, *ENSG00000308766*, *SCN3A*, and *TEKT2*, had the largest logFC differences between fetal sex (FDR < 0.05, ∣log2FC∣>1; Table 4). Overrepresentation analysis using ErmineJ yielded no statistically significant Gene Ontology (GO) biological processes across any lipid-candidate or genome-wide analysis group, indicating that no single pathway is highly represented or altered by our DEGs.

**Table 4:** DEGs unique to fetal female or male PE placentas.

| Gene name | Ensembl ID | Status | logFC<br>Females<br>(control vs<br>PE) | logFC<br>Males<br>(control vs<br>PE) |
| --- | --- | --- | --- | --- |
| **Lipid-Candidate Genes** |  |  |  |  |
| LDHB | ENSG00000111716.14 | Male-specific downregulation<br>(no difference in females) | - | -0.94253 |
| IGKC | ENSG00000211592.8 | Male-specific downregulation<br>(no difference in females) | - | -3.13703 |
| **Genome-wide** |  |  |  |  |
| NES | ENSG00000132688.11 | Female upregulation, Male<br>downregulation | 0.53526417 | - |
| STRIP2 | ENSG00000128578.10 | Female-specific<br>downregulation (no difference<br>in males) | -1.03078841 | - |
| ENSG0000<br>0296559 | ENSG00000296559.1 | Female-specific<br>downregulation (no difference<br>in males) | -0.56035345 | - |
| ENSG0000<br>0304336 | ENSG00000304336.1 | Female-specific<br>downregulation (no difference<br>in males) | -0.75789110 | - |
| ENSG0000<br>0224417 | ENSG00000224417.2 | Female-specific<br>downregulation (no difference<br>in males) | -1.07732579 | - |
| SYT16 | ENSG00000139973.18 | Female-specific<br>downregulation (no difference<br>in males) | -0.79985267 | - |
| C16orf54 | ENSG00000185905.4 | Female-specific<br>downregulation (no difference<br>in males) | -1.03301924 | - |
| ENSG0000<br>0308766 | ENSG00000308766.1 | Female-specific upregulation<br>(no difference in males) | 4.16608618 | - |
| ENSG0000<br>0280247 | ENSG00000280247.1 | Female-specific upregulation<br>(no difference in males) | 0.83798275 | - |
| ENSG0000<br>0260990 | ENSG00000260990.2 | Female-specific upregulation<br>(no difference in males) | 1.10111446 | - |
| TAS2R15P | ENSG00000212125.2 | Female-specific upregulation<br>(no difference in males) | 1.04875014 | - |
| SCN7A | ENSG00000136546.17 | Female-specific upregulation<br>(no difference in males) | 0.67667815 | - |
| IGKC | ENSG00000211592.8 | Male-specific downregulation<br>(no difference in females) | - | -3.28525 |
| SCN3A | ENSG00000153253.20 | Male-specific downregulation<br>(no difference in females) | - | -3.70501 |
| CDH13 | ENSG00000140945.17 | Male-specific downregulation<br>(no difference in females) | - | -1.43219 |
| ENSG00000292977 | ENSG00000292977.2 | Male-specific downregulation (no difference in females) | - | -1.41048 |
| LDHB | ENSG00000111716.14 | Male-specific downregulation (no difference in females) | - | -1.01704 |
| SLC13A4 | ENSG00000164707.17 | Male-specific downregulation (no difference in females) | - | -0.99318 |
| PKD1L1 | ENSG00000158683.10 | Male-specific upregulation (no difference in females) | - | 1.197248 |
| ENSG00000298360 | ENSG00000298360.1 | Male-specific upregulation (no difference in females) | - | 1.636191 |
| TGFBR3L | ENSG00000260001.8 | Male-specific upregulation (no difference in females) | - | 1.215877 |
| ENSG00000300349 | ENSG00000300349.1 | Male-specific upregulation (no difference in females) | - | 1.124787 |
| ENSG00000286339 | ENSG00000286339.2 | Male-specific upregulation (no difference in females) | - | 1.883385 |
| IGKC | ENSG00000211592.8 | Male-specific upregulation (no difference in females) | - | 0.77558 |
| ENSG00000305011 | ENSG00000305011.1 | Upregulated in both females and males (males to greater extent) | 0.87471373 | 3.172599 |

### 3.3 RNA Microarray Validation Cohort Reinforces Sex-Specific Differential Expression in PE

To validate our RNA-seq findings in an independent cohort, we assessed differential gene across our significant hits using an independent Affymetrix Human Gene 1.0 microarray dataset. In the lipid- candidate genes, 51 of 66 autosomal genes successfully validated in the combined-sex population (77.3% validation). Sex-stratification confirmed 37 of 58 autosomal genes validated in the female population (63.8% validation), and 61 of 78 validated in the male population (78.2% validation; Supplementary Table 4).

Genome-wide validation confirmed 168 of 229 autosomal genes in the combined-sex group (73.4% validation), 104 of 192 in the female-stratified (54.2% validation), 178 of 299 in the male-stratified (59.5% validation; Supplementary Table 5).

### 3.4 Targeted lipid-candidate and genome-wide analyses reveal differential isoform expression masked by differential gene expression analyses in PE

To evaluate transcript-level variations and preferential isoform expression in PE, we performed differential isoform expression analysis using Swish across both our targeted lipid-candidate gene set and the whole genome. In the combined-sex lipid-candidate analysis, Swish identified 103 significantly upregulated isoforms across 57 genes and 34 downregulated isoforms across 29 genes (FDR<0.05,∣log2 FC∣>1). Interestingly, DESeq2 flagged only a fraction of these isoform-level changes as DEGs (30 upregulated, 6 downregulated), demonstrating that standard DEG analyses mask changes in isoform activity (Figure 4). The strongest transcript shifts included isoform downregulations in *ACOXL* and *PRRX1*, and isoform upregulations in *LEP* and *FLT1* (Supplementary Table 6).

**Figure 4:**
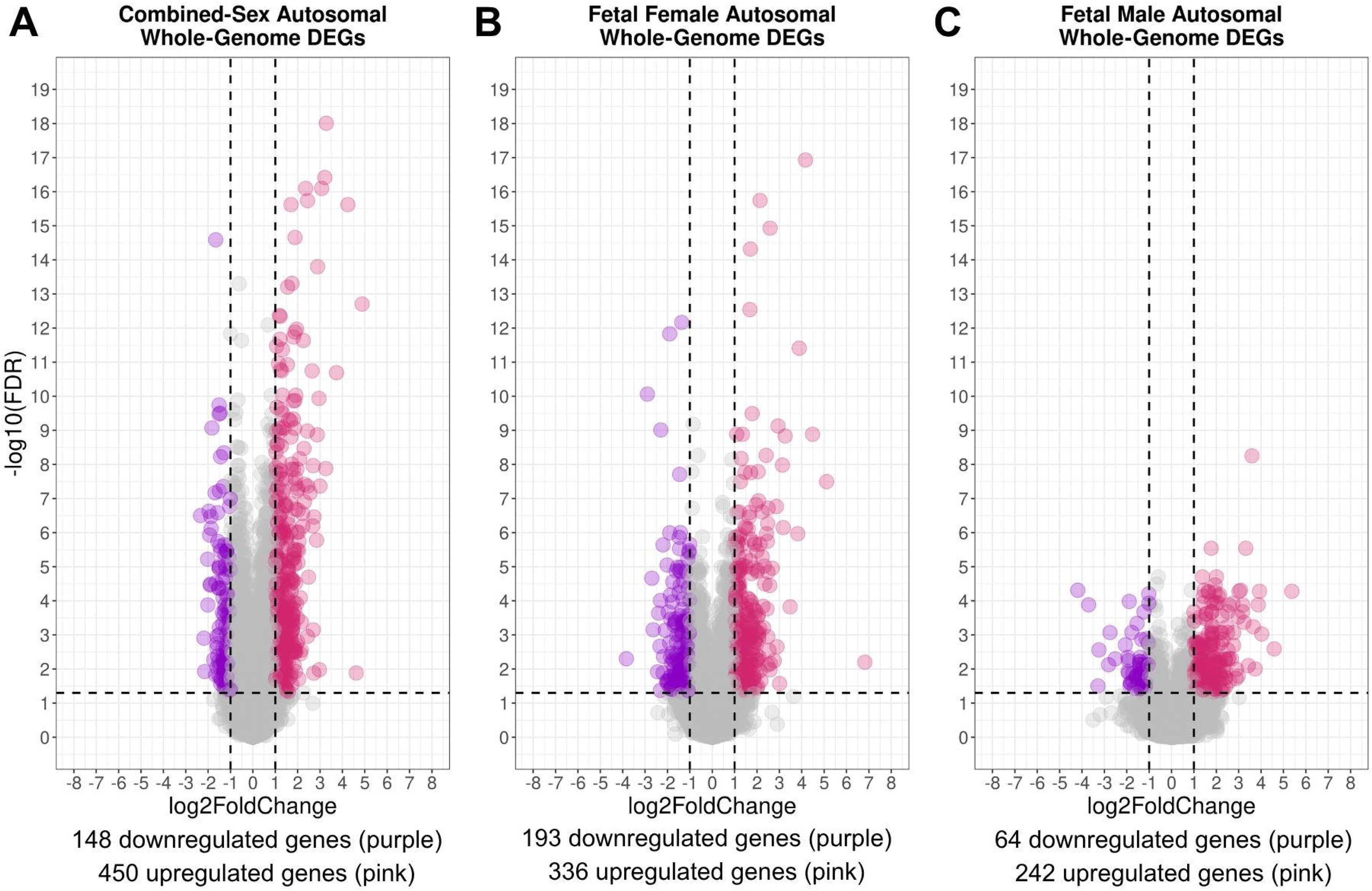
Scatter plots comparing the overlap between differential gene (DESeq2) and isoform (Swish) expression analysis. Each data point represents a single gene isoform tested for both differential gene and isoform expression, with the color indicating whether both methods yield consistent results. Grey points represent genes that are non-significant in both analyses, blue indicates genes significant only in the gene-level analysis (DESeq2), green indicates genes significant only at the isoform level (Swish), and red highlights genes that are significant in both analyses. The top row reflects analyses in lipid-candidate genes, while the bottom row reflects whole-genome analyses. The columns represent the combined fetal-sex, female-only, and male-only analysis groups.

Sex-stratified lipid-gene analysis showed that females placentas exhibited 83 upregulated isoforms across 48 genes (27 designated as DEGs) and 49 downregulated isoforms across 42 genes (12 DEGs). Male lipid analysis yielded fewer changes, with 39 upregulated isoforms across 27 genes (18 DEGs) and 11 downregulated isoforms across 11 genes (3 DEGs) (Supplementary Table 6). Among these analyses, only 22 differentially expressed isoforms were shared between sexes, all with matching up- and downregulation.

Genome-wide analysis in the combined-sex population identified 420 significantly upregulated isoforms across 193 genes (86 of which were DEGs) and 127 downregulated isoforms across 100 genes (29 DEGs) in PE placentas relative to controls (Supplementary Table 6). In the sex-stratified genome-wide analyses, female placentas displayed 316 upregulated isoforms across 163 genes (66 DEGs) and 155 downregulated isoforms across 118 genes (34 DEGs) in PE (Supplementary Table 6). Male placentas exhibited fewer changes again, showing 109 upregulated isoforms across 77 genes (37 DEGs) and 36 downregulated isoforms across 32 genes (13 DEGs; Supplementary Table 6). Sixty-seven differentially expressed isoforms were shared between sexes, all with matching up- and downregulation.

### 3.5 Alternative splicing analysis identifies several mutually exclusive exon and skipped exon events in PE placentas

To determine the splicing patterns driving these shifts in isoform abundance, we utilized rMATS to screen for alternative splicing events in PE. The software identifies five basic alternative splicing events: 1) skipped exon, where a specific exon is excluded from the mature mRNA; 2) alternative 5’ splice site, the utilization of a different splice site at the 5’ end; 3) alternative 3’ splice sites, the utilization of a different splice site at the 3’ end; 4) mutually exclusive exons, where two neighboring alternative exons are situated between flanking exons, and the splicing machinery selectively includes one of the two in the mature mRNA; and 5) retained intron, where an intron is inappropriately kept in the mature mRNA rather than being spliced out [99].

In the combined-sex population, rMATS identified a total of 1785 significant differential splicing events across 1320 genes in PE placentas compared to controls. Mutually exclusive exons were the most prevalent alternative splicing patterns, accounting for 1478 events. Skipped exons represented the second most frequent modification with 154 significant events, followed by RIs at 83 events. Alternative 3’ splice sites and alternative 5’ splice sites occurred less frequently, contributing 43 and 27 significant events, respectively. These events physically alter transcript composition and may yield altered placental protein variants in PE.

### 3.6 Small fraction of lipid-candidate and genome-wide gene expression changes are driven by transcript-level and alternative splicing differences in PE

To investigate the upstream mechanisms driving the changes in gene- and transcript-level expression, we intersected our DESeq2, Swish, and rMATS datasets. Due to reduced statistical power in sex-stratified groups, we restricted rMATS mapping to the combined-sex whole-genome population. Across all tested groups, only a fraction (10% to 50%) of our identified DEGs exhibited corresponding differential isoform expression, with an even smaller fraction of those shifts stemming from underlying alternative splicing events. The remaining 50% to 90% of DEGs that did not feature differential isoform expression likely reflects the higher statistical thresholds required for isoform-level analysis, where evaluating multiple individual transcripts per gene increases number of hypothesis tests, and the rate of false negatives due to more stringent multiple-testing corrections [125].

In the combined-fetal-sex lipid-candidate analysis, nearly half of the detected DEGs (36 of 76) were driven by differential isoform expression. Specifically, 12 of 76 DEGs were driven by expression changes across all annotated isoforms, 13 were driven by expression changes in a single isoform, and 11 were driven by expression changes across multiple distinct isoforms (Figure 5). *ACOXL* was the only target where gene-level alterations directly involved an underlying alternative splicing event, specifically a mutually exclusive exon switch. In the female lipid analysis, 36 of 75 DEGs were driven by isoform-level shifts, where 14 were driven by expression changes across all isoforms, 11 were driven by changes in a single isoform, and 11 were driven by changes in more than one isoform (Figure 5). In the male lipid analysis, 23 of the 97 DEGs were driven by differentially expressed isoforms, 7 were driven by expression changes across all isoforms, 10 were driven by changes in a single isoform, and 6 were driven by changes in multiple isoforms (Figure 5).

**Figure 5:**
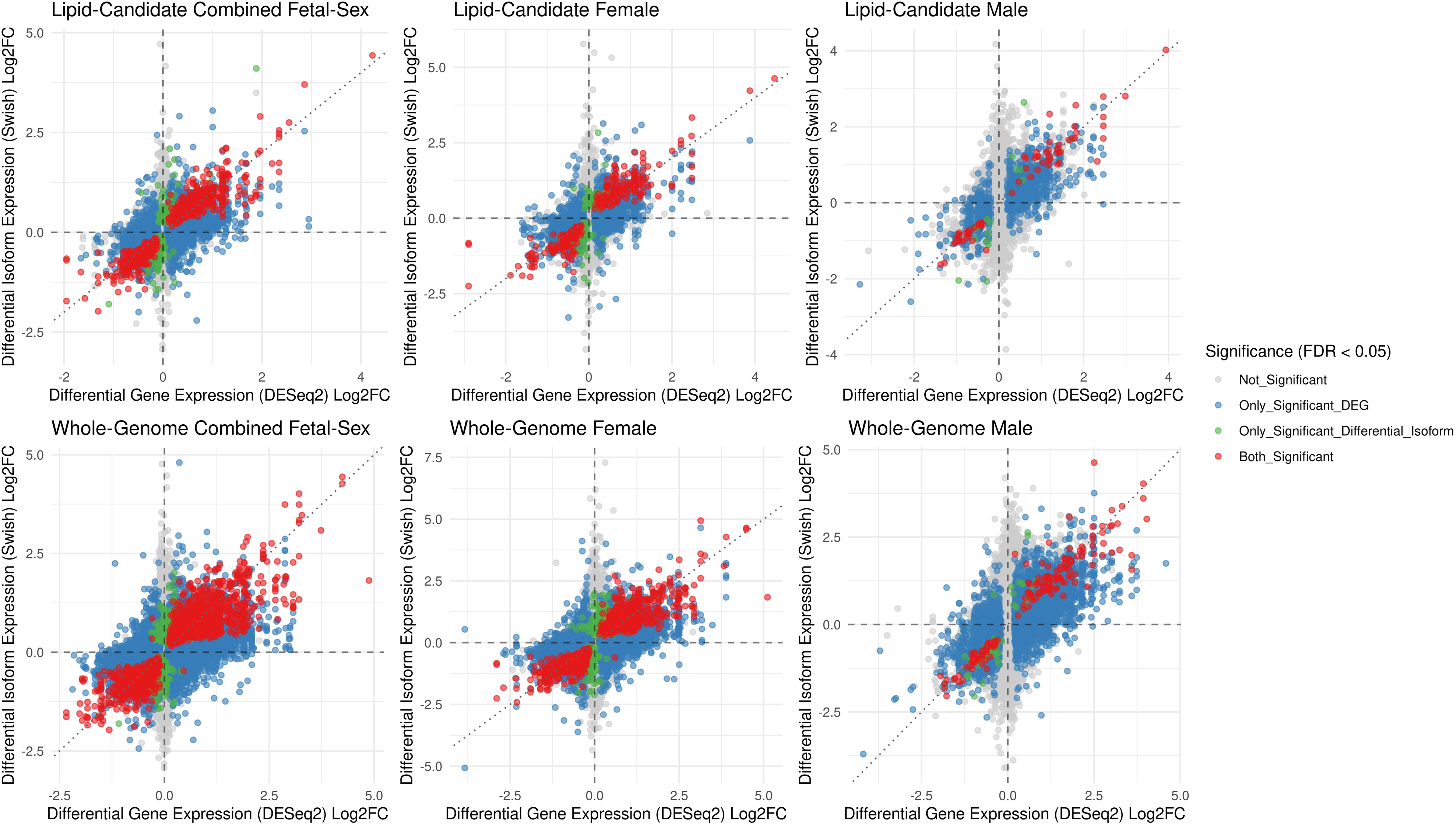
Bar plot illustrating the percentage of DEGs across the six experimental groups categorized by their specific differential transcript expression (DTE) profiles. Categories indicate whether a gene’s expression change is driven by a single isoform, several isoforms, all isoforms, or features no significant individual transcript changes. X-axis labels refer to the combined fetal-sex, female-only, and male-only analysis groups (combined, F, M) for the lipid-candidate gene (Lip) and whole-genome (WG) analyses.

Genome-wide, only 151 of 598 total DEGs in the combined-sex group involved individual isoform shifts. Within these targets, 41 DEGs were driven by expression changes across all annotated isoforms, 63 were driven by expression changes in a single isoform, and 47 were driven by expression changes across multiple isoforms (Figure 5). Overall, only five alternative splicing events drove gene-level changes. When stratified by sex, female placentas featured 132 transcript-driven DEGs out of 508 total targets, comprising 40 driven by expression changes across all isoforms, 46 driven by a single isoform, and 46 driven by multiple isoforms. Male placentas featured only 74 transcript-driven DEGs out of 633 total targets, comprising 16 driven by expression changes across all isoforms, 39 driven by a single isoform, and 19 driven by multiple isoforms (Figure 5).

### 3.7 Differential isoform usage in PE is present, sex-specific, and undetected by differential gene expression analysis

To identify whether shifts in isoform usage undetected by DESeq2 may contribute to PE pathology, we ran IsoformSwitchAnalyzeR within the lipid-candidate genes and across the whole genome [101]. The software detects reciprocal shifts in preferential isoform usage between conditions and predicts their structural and functional consequences on protein-coding capacity, open reading frames (ORFs), intrinsically disordered regions (IDRs), signal peptides, or functional domains. These changes can prevent protein production, modify binding capacity, shift translation start sites, or change protein localization, potentially driving functional gains or losses [101,106,126–128].

In the lipid-candidate combined-sex analysis, 5 genes exhibited significant isoform switches (Q < 0.05; Table 5). Functional consequence modeling predicted that the preferred PE isoform of *RHOF* gained an ORF elongation and functional domain, whereas the preferred PE isoform of *IGHG1* underwent ORF shortening. Interestingly, *IGHG1* was the only gene with an isoform switch that also showed differential gene expression in DESeq2, highlighting that standard differential gene expression analysis may miss switches in isoform usage.

**Table 5:**
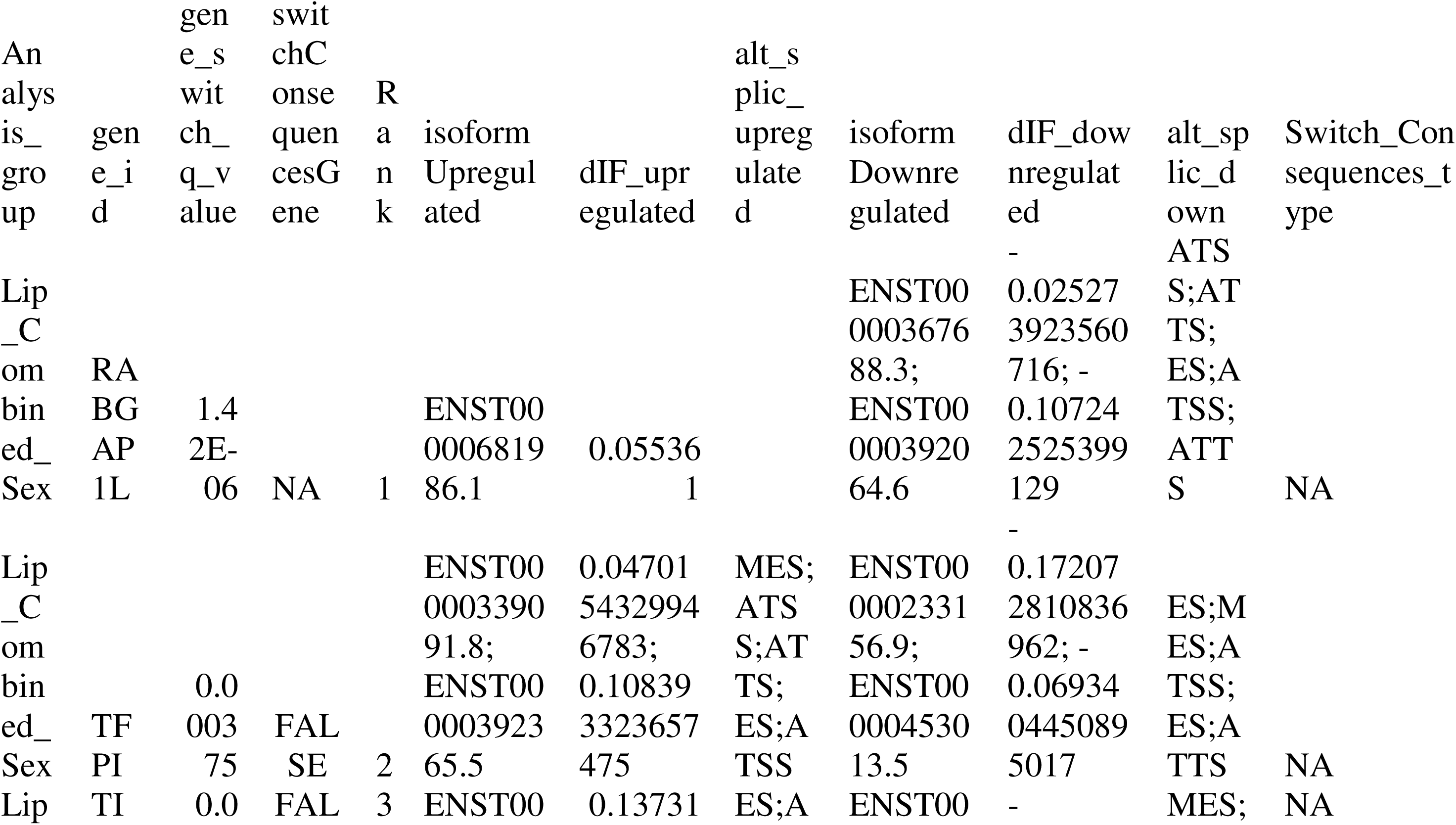
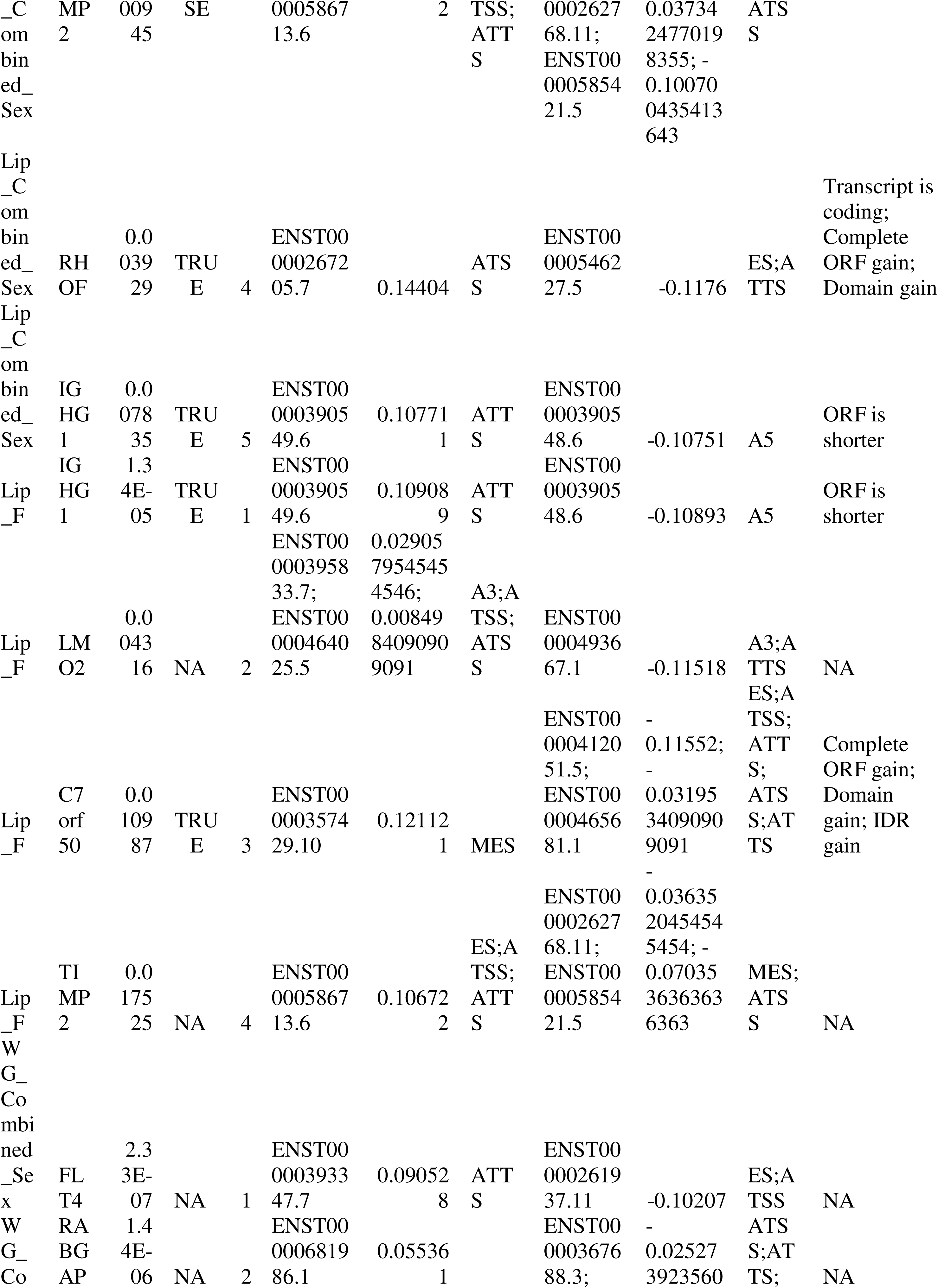
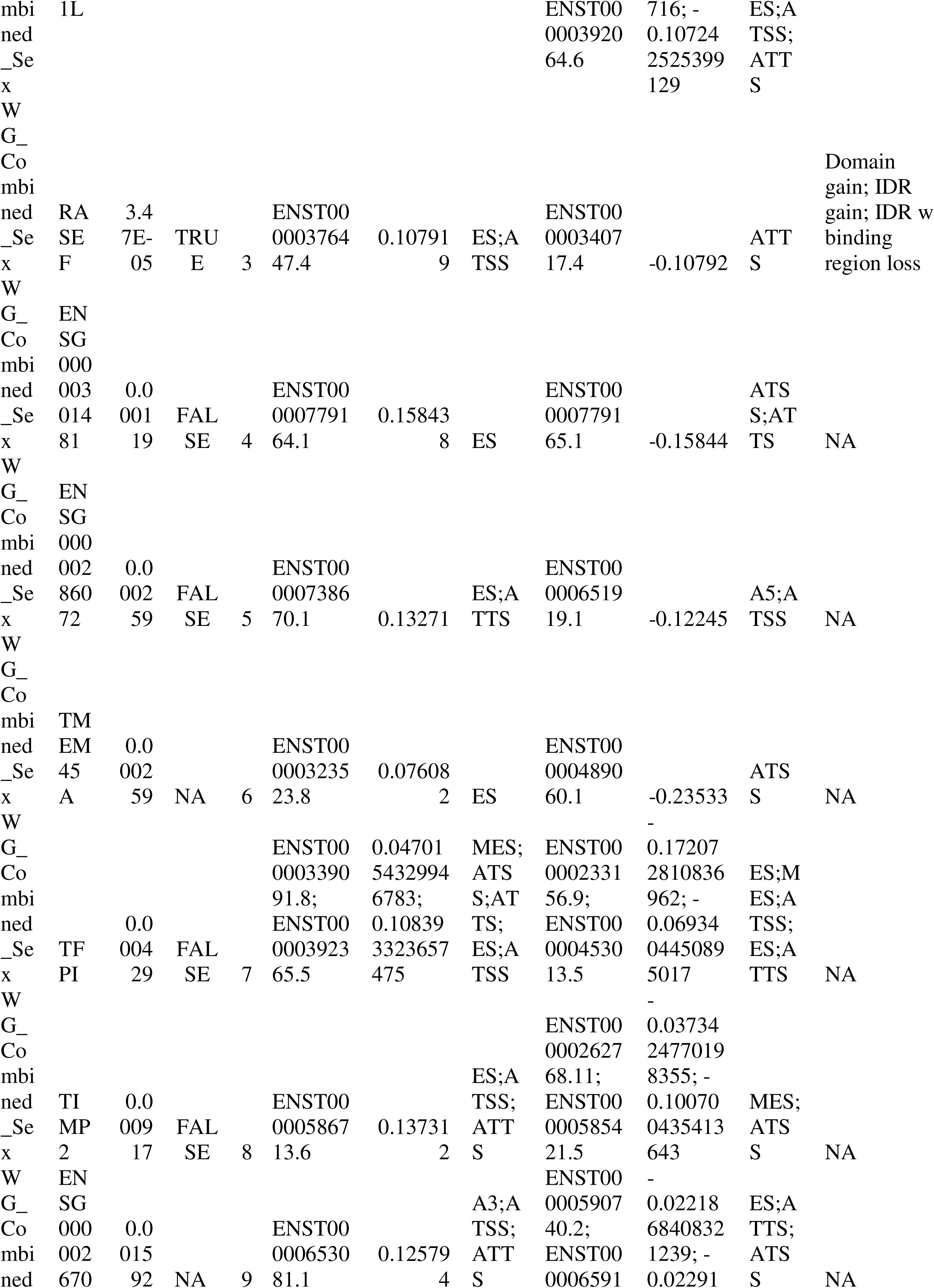
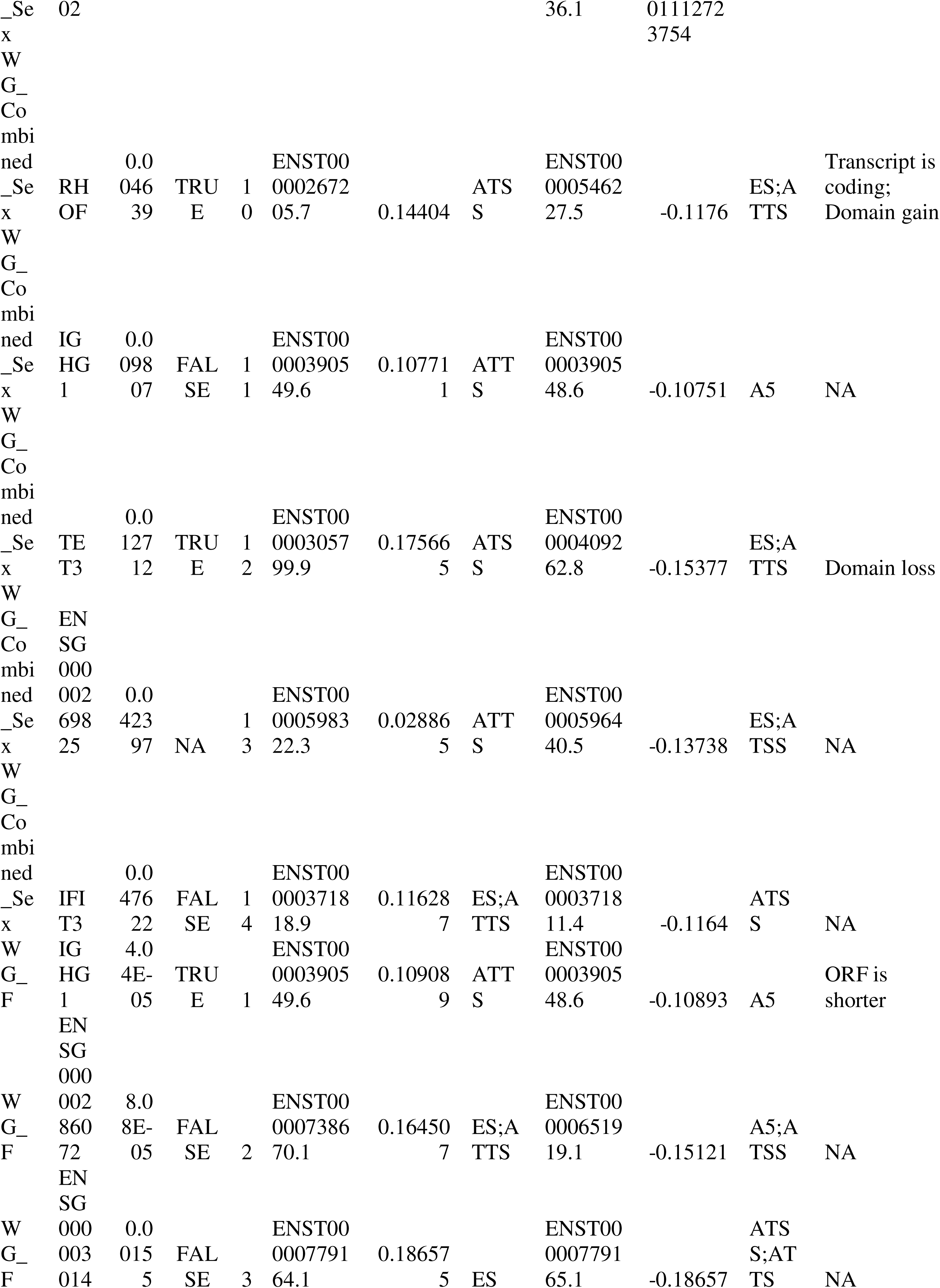
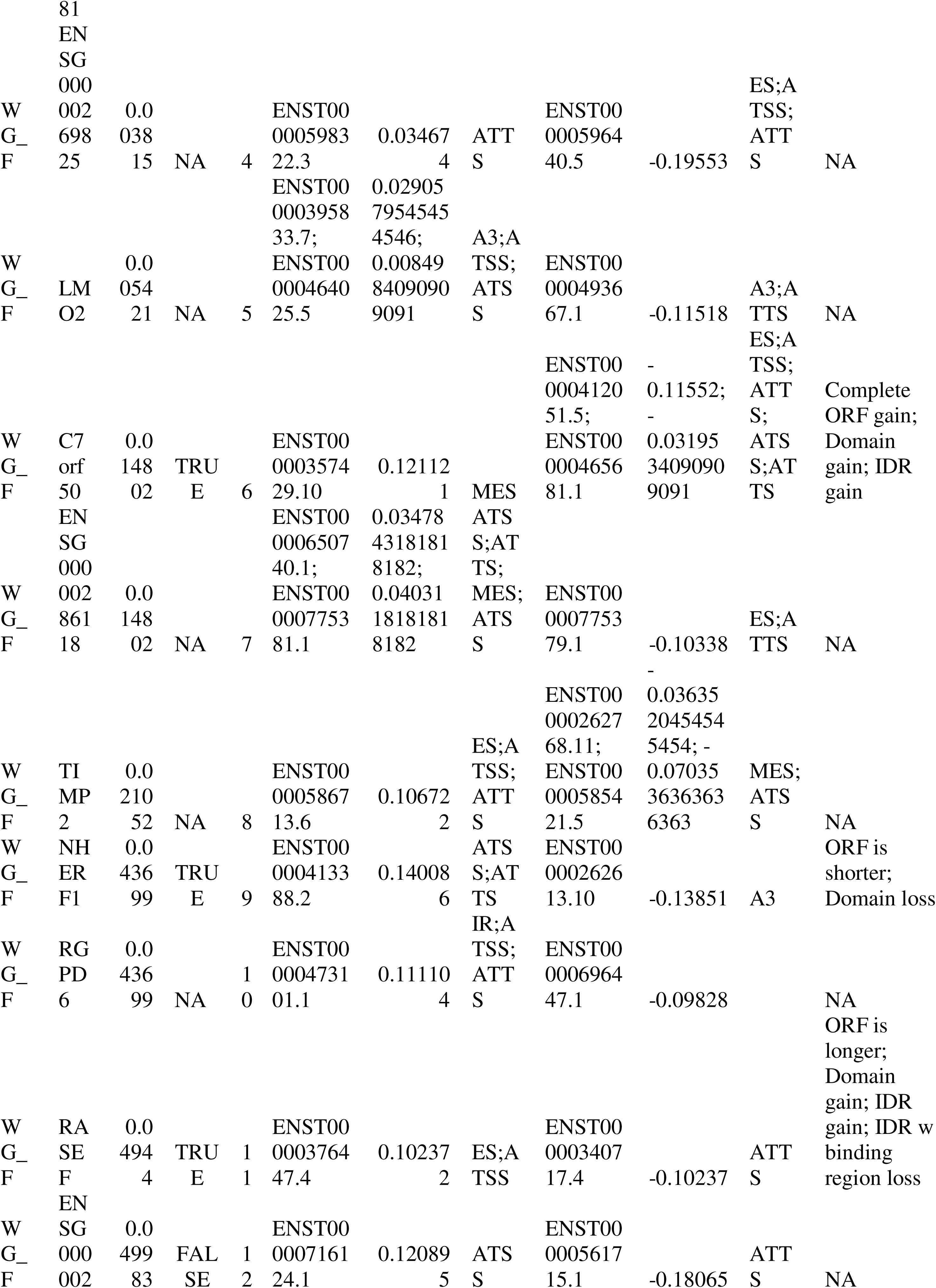
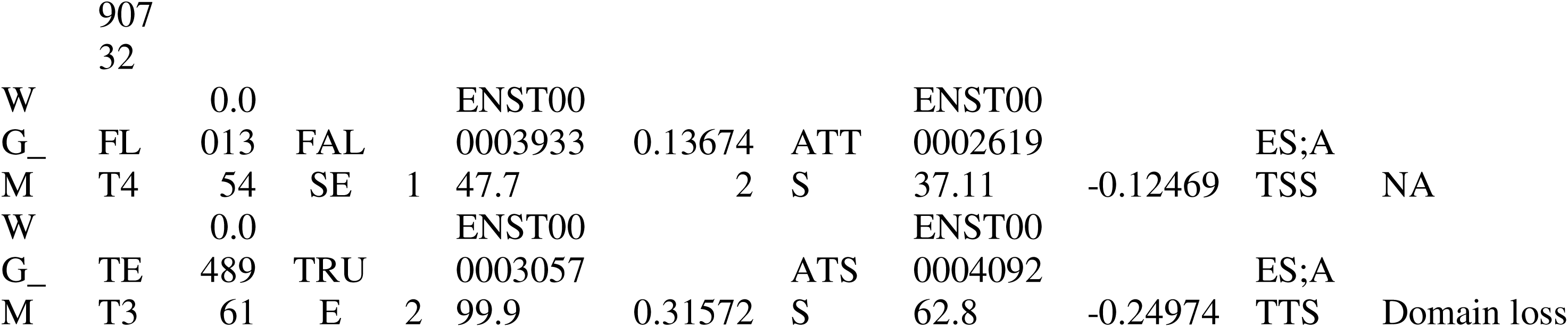
List of genes undergoing isoform switching in PE placentas (IsoformSwitchAnalyzeR).

**Table 6:**
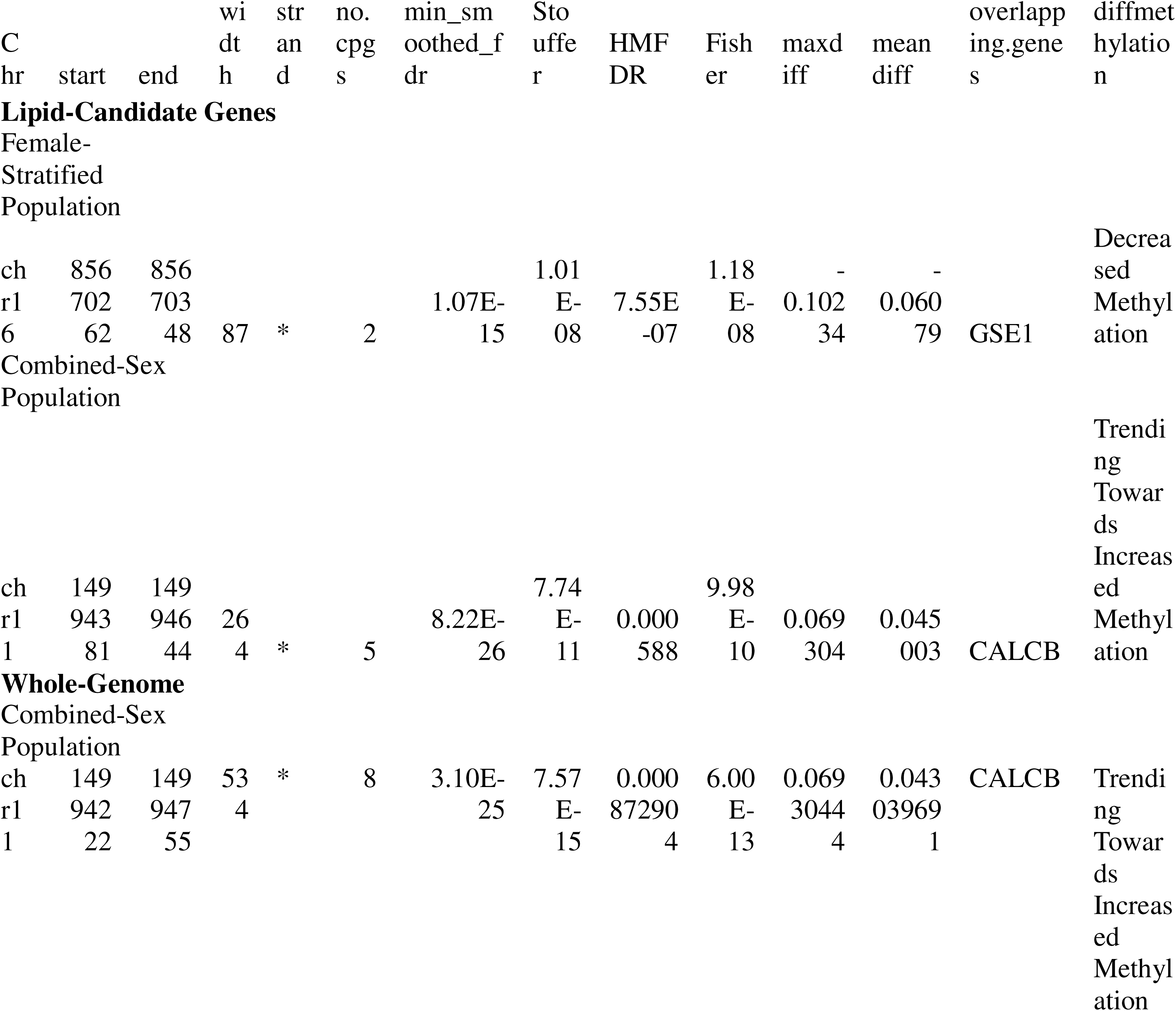
List of differentially methylated regions (DMRs) across genome in preeclampsia.

After sex stratification, the female lipid analysis revealed 4 major isoform switches (*TIMP2*, *IGHG1*, *LMO2*, and *C7orf50*), none of which were flagged as DEGs in DESeq2. Fetal males exhibited no significant isoform switches within the lipid-candidate analysis, suggesting that the combined-sex signals were driven primarily by female-specific dynamics.

Genome-wide analyses identified 14 significant isoform switches across the combined-sex group (Q < 0.05; Table 5), with only 5 overlapping with DEGs. Sex stratification identified 12 significant isoform switches in the female-stratified genome, 4 of which carried predicted functional and structural consequences (Table 5). The male genome-wide analysis identified 2 significant isoform switches (*FLT4* and *TET3*), with only *TET3* overlapping with a DEG and undergoing a predicted structural domain loss. Interestingly, across all analyses, the majority of these isoform switches were driven by alternative transcription initiation and termination sites, rather than conventional alternative splicing events like exon skipping or mutually exclusive exons.

### 3.8 No differentially methylated CpGs in lipid-candidate genes and genome-wide between PE and control pregnancies

To determine whether the changes in transcriptional and alternative splicing patterns were caused by DNAm changes, we performed a separate meta-analysis across four public placental Illumina 450K DNA methylation datasets from PE and control placentas.

Statistically significant differentially methylated CpGs were defined by an FDR < 0.05 and a ∣Δβ∣ > 0.05 to indicate potential biological significance. Across both the targeted lipid-candidate and the whole-genome analyses, no CpGs reached statistical significance between control and PE placentas in combined-sex or sex-stratified interaction term models (Figures 6a-c, 7a-c). Similarly, no differential DNAm was observed on the X chromosome in either sex (Supplementary Figures 7, 8), and the interaction model identified no sex-specific epigenetic differences. ErmineJ analysis yielded no significantly overrepresented biological processes across any candidate or genome-wide DNA methylation group.

**Figure 6:**
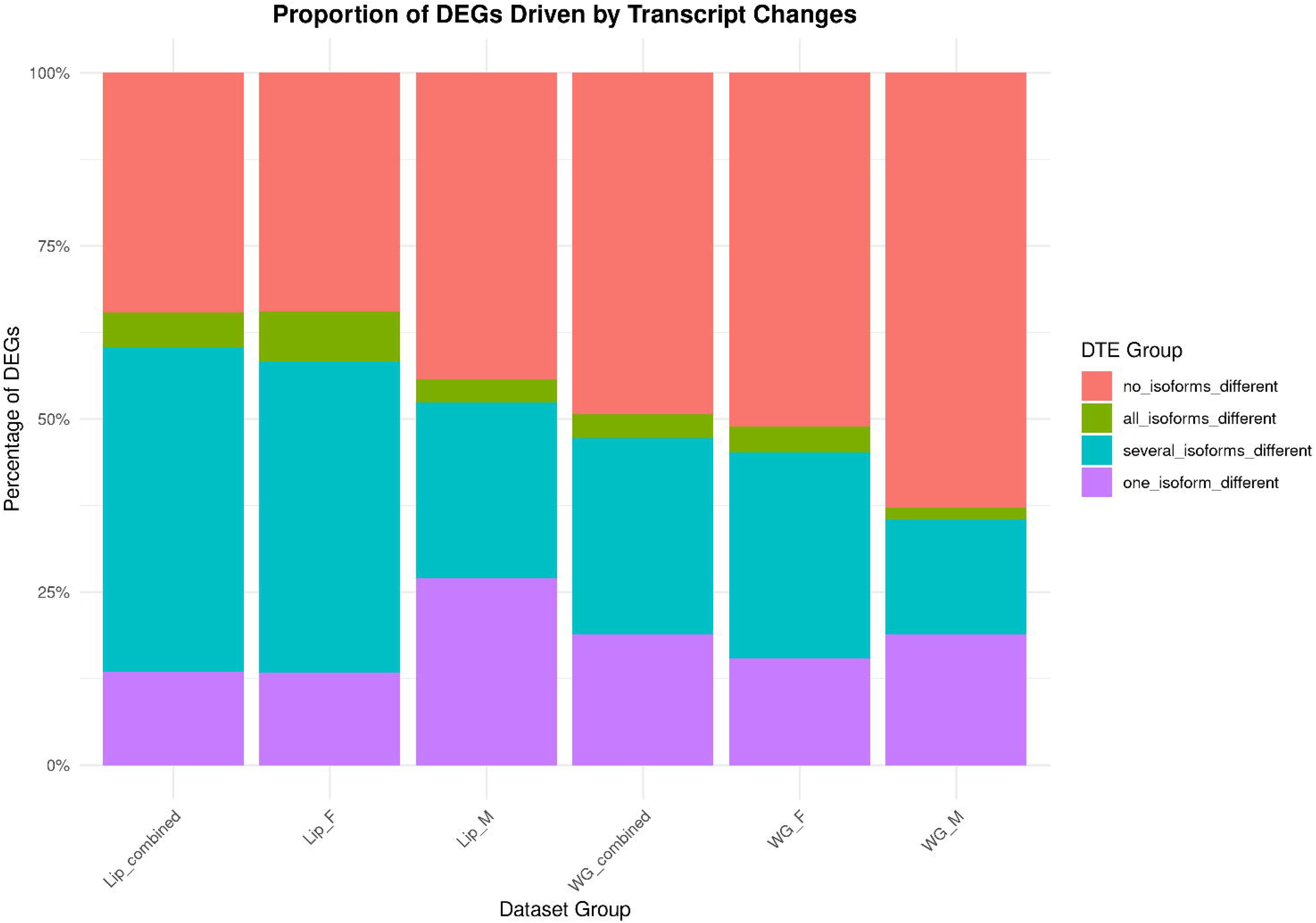
Volcano plot depicting changes in autosomal DNAm in lipid-candidate genes between PE and control placentae in the (A) combined fetal sex population (n = 172; 100 PE, 72 Control), (B) female-stratified population (n = 83; 48 PE, 35 Control), and (C) male-stratified population (n = 89; 52 PE, 37 Control).

**Figure 7:**
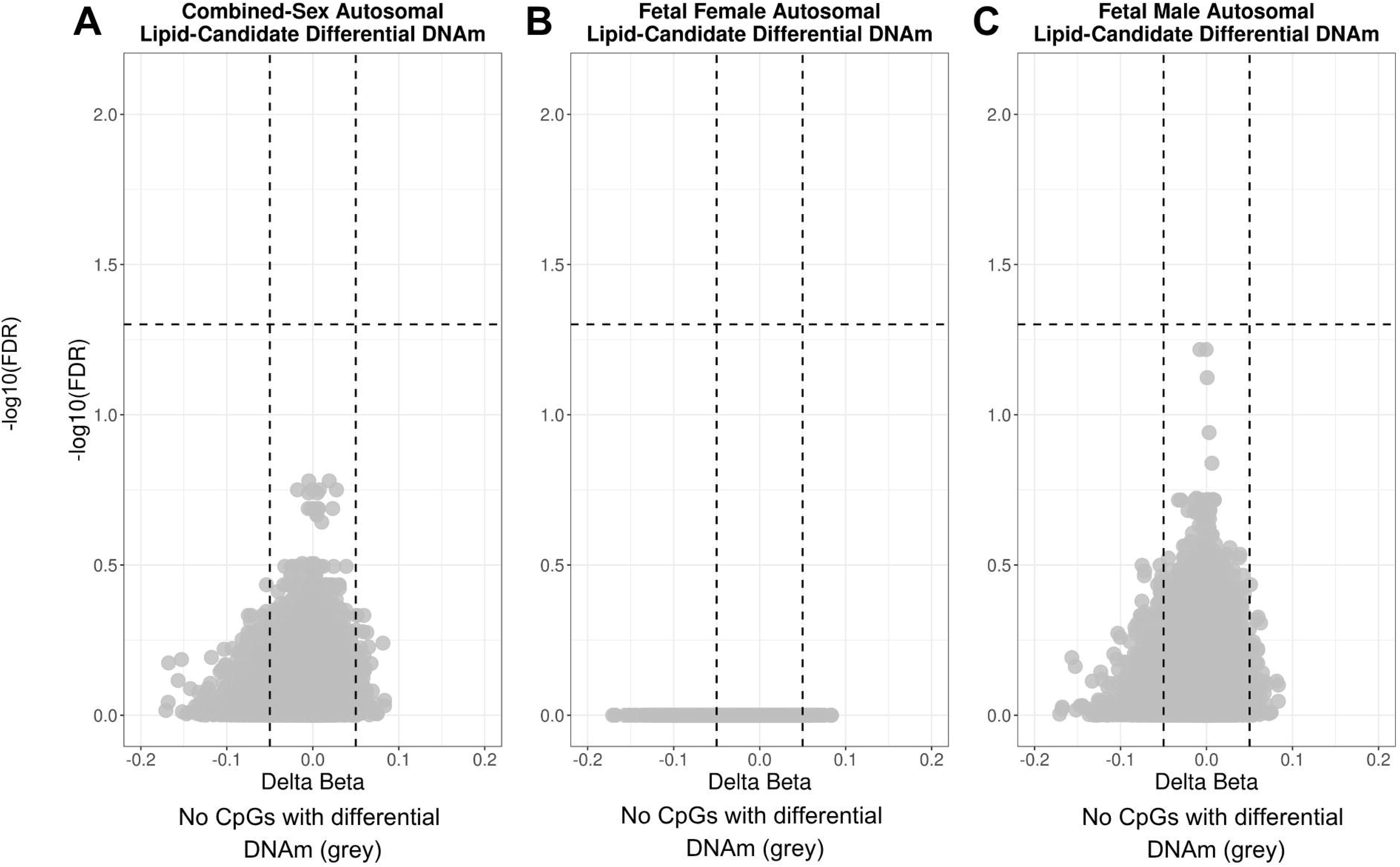
Volcano plot depicting changes in autosomal DNAm genome-wide between PE and control placentae in the (A) combined fetal sex population (n = 172; 100 PE, 72 Control), (B) female-stratified population (n = 83; 48 PE, 35 Control), and (C) male-stratified population (n = 89; 52 PE, 37 Control).

### 3.9 Fetal female placentas revealed biologically significant differentially methylated region (DMR) in *GSE1* and *CALCB* in the combined-sex population

To identify broader epigenetic shifts beyond individual CpGs, we performed DMR analysis on lipid- candidate genes and across the genome. In the lipid-candidate gene analysis, 1 significant DMR showed decreased regional DNAm within the female-stratified population (FDR < 0.05, mean Δβ<−0.05; Table 5), mapping to the *GSE1* gene. No biologically significant DMRs were identified in the combined-fetal-sex or male-stratified populations, however, one DMR in the *CALCB* gene, trended towards increased methylation (FDR < 0.05, mean Δβ > 0.00; Table 5) in the combined-sex population. Expanding this analysis to the whole genome yielded no significant DMRs across the sex-stratified conditions, but the same region in the *CALCB* gene showed increased regional methylation in the combined-sex population (Table 5).

### 3.10 Placental cell-type proportions differed between sex and conception type in DNAm and RNA-seq

To assess whether the altered epigenetic and expression patterns were driven by shifting cell-type proportions in PE, we performed cell-type deconvolution analyses using both our RNA-seq and DNAm data. In the RNA expression cell deconvolution results, female preeclamptic placentas showed increased proportions of fetal mesenchymal stem cells and fetal nucleated red blood cells (nRBCs) relative to healthy control placentas (Figure 8a). We observed no other proportional differences between control and PE placentas. However, baseline sex-differences were observed between healthy controls, with male placentas showing higher proportions of fetal endothelial cells and fetal fibroblasts than female controls. Male PE placentas showed an increased proportion of fetal GZMK natural killer cells compared to female PE placentas (Figure 8a).

**Figure 8:**
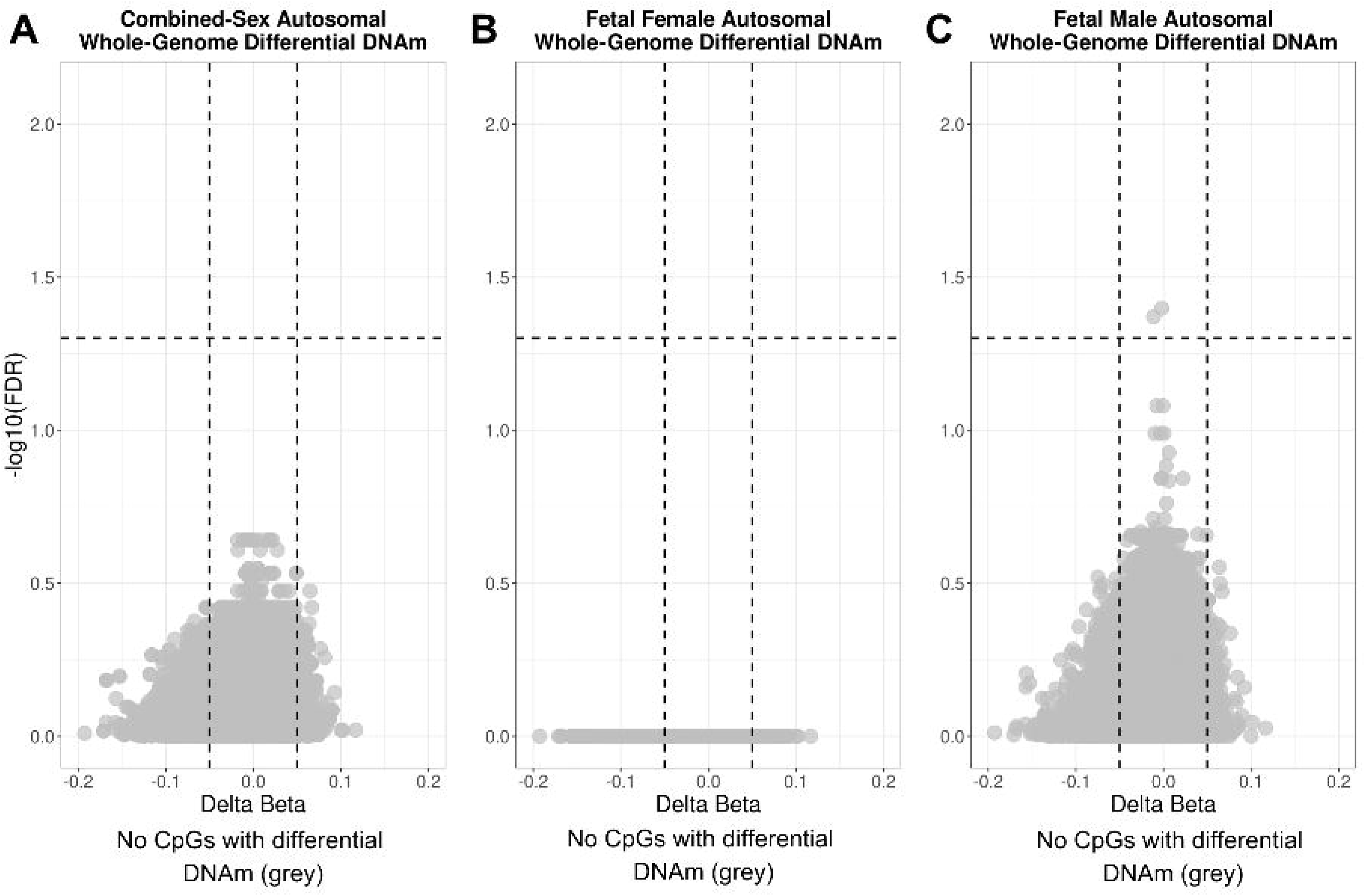
Placental cell-deconvolution analyses performed using (A) RNA-seq data and (B) DNAm data. Groups are stratified by fetal sex and disease group: female controls (CONT_F), male controls (CONT_M), female preeclampsia (PE_F), male preeclampsia (PE_M).

Cell deconvolution of the DNA methylation array partially diverged from the RNA-seq-based results. This divergence is likely due to lower cell-type resolution in the DNAm model, preventing more specific cell type abundances from being extrapolated. The DNAm data showed a significantly higher abundance of stromal cells in both the male and female PE placentas compared to controls (Figure 8b). Syncytiotrophoblasts showed a higher abundance in PE males, and trophoblasts showed a higher abundance in PE females compared to the controls of either sex (Figure 8b). However, the DNAm data revealed no significant differences in the abundance of fetal nucleated red blood cells, mesenchymal stem cells, or endothelial cells.

## 4. Discussion

This study aimed to investigate how lipid pathways are regulated in PE, and infer downstream molecular implications of this regulation using full term placental RNA sequencing and DNAm data. We first restricted RNA-seq data to genes implicated in lipid-related pathways throughout the literature. This candidate-gene approach lowered the number of gene comparisons between groups to improve statistical sensitivity and capture more subtle changes in lipid-specific genes [84].

### Lipid-related genes have altered expression in PE

In the combined fetal-sex population, our candidate analysis identified 76 DEGs (FDR < 0.05, ∣log2 FC∣>1) between control and PE placentas. *LEP*, *FLT1*, *ARMS2*, *HSD3B2*, and *IGHG2* were among the top DEGs by log-fold change, aligned with previous transcriptomic studies that highlight disruptions in lipid processing genes in PE pregnancies [129,130].

We then broadened our analysis to an un-filtered genome-wide differential expression analysis. In the combined fetal-sex population there were 598 DEGs between PE and controls, with top genes including *LEP* once again, alongside *HTRA4*, *PRRX1*, and two uncharacterized long non-coding RNAs (lncRNAs), *ENSG00000296461* and *ENSG00000302259*. Many studies have identified altered *LEP* expression in PE [69,131,132]; the gene encodes a hormonal protein that regulates key pathways implicated in PE pathology including glucolipid metabolism, angiogenesis, blood pressure, immune response, and intrauterine fetal development [5,133]. Other DEGs similarly participate in key aspects of PE pathology, including steroid biosynthesis [134], humoral immunity [135], and extravillous trophoblast migration [136,137]. The functional diversity of these genes highlights the heterogeneity in pathway dysregulation in PE placentas.

We performed an expression validation cohort using an independent Affymetrix microarray dataset to validate the differential expression patterns identified via RNA-seq. Overall, the candidate lipid genes validated at a higher rate (77.3%) than the genome-wide genes (73.1%) in the independent dataset. The high reproducibility of these differential gene expression patterns, particularly among lipid-related identified in the genome-wide analyses, underscores the potential clinical utility of lipid-candidate genes for future predictive or diagnostic tools.

Interestingly, despite this cross-platform validation, functional enrichment analysis via ErmineJ yielded no significant overrepresented Gene Ontology (GO) pathways across any of the analysis groups. This finding suggests that PE is a heterogeneous disorder that impacts multiple organ systems and alters several lipid pathways, rather than being driven by a single dominant one.

### PE transcriptome is altered depending on fetal sex

Previous reports have indicated that analyzing male and female placentas together as a single population may mask fetal-sex differences in PE [138]. To address this concern, we stratified by sex in our analyses. In the sex-stratified lipid-candidate analysis, we identified 73 autosomal DEGs in females and 58 in males. In the whole-genome analysis, we identified 529 DEGs in females and 306 in males (FDR < 0.05, ∣log2FC∣>1).

We then reanalyzed the lipid-candidate and whole-genome datasets using a sex-interaction term model to confirm that observed sex-specific transcriptomic shifts reflected true biological changes rather than sampling artifacts or power loss from stratification [83]. While sex-stratification analyzes male and female cohorts independently, the interaction term model calculates baseline variation across all samples, increasing power and allowing us to evaluate whether differential gene expression in PE differs by sex [139]. For females, DEG numbers and gene identities were nearly identical between the stratified and interaction analyses. For males, however, the interaction model identified more than double the DEGs found in the sex-stratified analysis. This suggests that the interaction model reduces false negatives in males by estimating shared background variance in all samples, increasing statistical power for DEG discovery in the smaller male cohort [140]. These findings highlight that interaction terms provide a strong alternative model for identifying sex-specific pathological mechanisms in unbalanced cohorts, without sacrificing statistical power.

Interestingly, in our sex-stratified analyses, we observed several DEGs unique to either females or males that were also confirmed as sex-specific in our validation cohort. To further explore this sex-specific impact of PE and ensure these differences were not driven by sampling artifacts between sexes, we further extrapolated the results from the interaction term model to identify whether PE differentially affects gene expression based on fetal sex [83]. The targeted lipid-candidate analysis highlighted two genes with sex-specific expression patterns, *LDHB* and *IGKC*. Both genes showed significant downregulation in male PE placentas but no differential expression in female PE placentas. *LDHB* encodes lactate dehydrogenase B, an enzyme that converts lactate to pyruvate for aerobic metabolism [141]. This downregulation could indicate that male PE placentas might have a reduced capacity to consume pyruvate for oxidative metabolism, leaning more towards anaerobic glycolysis for ATP production compared to female PE placentas. The *IGKC* gene encodes a structural component of antibodies that facilitates the clearance of bound antigens [142]. Its downregulation could therefore suggest that male PE placentas may exhibit a more suppressed humoral immune response compared to female PE placentas.

The whole-genome interaction term analysis further highlighted the functional diversity of genes showing differential regulation between male and female PE placentas. One DEGs with increased expression in exclusively in female PE placentas was *ENSG00000308766*, an antisense RNA to *FSTL3* (follistatin-like 3) [143]. High expression of the *FSTL3* glycoprotein is associated with the development of PE throughout gestation [144]. Increased expression of this antisense RNA in female PE placentas could alter *FSTL3* transcript abundance, potentially driving sex-specific differences in how *FSTL3* influences PE susceptibility.

Interestingly, 3 of the 11 genes downregulated only in male PE placentas, *LDHB*, *ACSS1*, and *DBI*, play a role in promoting oxidative glycolysis or lipid metabolism [141,145,146]. This finding reinforces the possibility that male PE placentas shift towards anaerobic metabolism to a greater extent than female PE placentas. The literature reports an increased glucose dependency in male trophoblasts in gestational diabetes [147] and higher expression of hypoxic markers in male PE placentas [148], aligning with our findings. Although one study observed an upregulation of fatty acid oxidation and uptake genes in male PE placentas [149], its focus on a small subset of genes may explain its discrepancy with our findings.

Also, many sex-specific DEGs mapped to uncharacterized non-coding genes (*ENSG00000298360*, *ENSG00000224417*, *ENSG00000296559*), highlighting a potential role for non-coding RNAs in driving sex-specific PE pathology [143]. Other sex-specific DEGs spanned diverse functions including cell structural components [150,151], trophoblast invasion [152], transcription factor regulation [153], vesicular trafficking [154], or immune system involvement [142].

### Alternate transcription initiation and termination sites drive isoform usage shifts in PE

We next investigated differential isoform expression to identify possible mechanisms driving differential gene expression in PE. These changes may stem from broad transcriptional shifts that alter all gene isoforms, or from the shift towards a specific preferred isoform via alternative splicing, initiation, or termination sites [38,39,155]. Our analyses revealed that PE transcriptomic changes are driven by a combination of the two mechanisms. For example, *LEP* and *FLT1* showed expression increases across all or most isoforms, whereas other DEGs, such as *LTF*, were driven by a single, preferentially expressed isoform. These findings highlight that PE-associated changes in gene expression result from both overall differential transcriptional activity and preferential isoform expression.

We then performed alternative splicing analysis in the whole-genome to determine whether the preferential expression of certain isoforms was driven by alternative splicing. Although we identified several differential splicing events (1,785 across 1,320 genes), only 5 of these events overlapped with the DEGs from DESeq2, and only 33 matched the differentially expressed isoforms from Swish. This finding indicates that alternative splicing is unlikely to he the primary driver of these preferentially expressed isoforms. Instead, alternative transcription initiation and termination sites may lead to these isoform differences [155,156].

In our differential isoform expression analysis, we identified several transcripts not designated as DEGs at the gene level. We thus sought to analyze isoform switching, reciprocal shifts in isoform expression within a gene. Most identified switches were undetected in differential gene expression analyses, highlighting that gene-level analyses alone can overlook underlying pathophysiological changes in gene structure. The predominant biological mechanisms underlying these switches were alternative transcription initiation or termination sites. For example, *FLT4* exhibited an isoform switch driven by an alternative transcription termination site, though no downstream functional protein changes were predicted. *FLT4* encodes VEGFR-3, a regulator of endothelial cell migration and angiogenesis [157]. Interestingly, this finding aligns with recent computational and experimental evidence demonstrating increased usage of alternative last exon 2 in *FLT4* transcripts from early-onset PE placentas compared to term controls [34]. This finding supports the low overlap between isoform switching and alternative splicing observed in our analysis, suggesting that alternative initiation and termination sites are the primary drivers of preferential isoform usage in PE. Future work should thus aim to characterize the roles of cleavage, polyadenylation, and transcription factors as drivers of these changes and PE pathology [158,159]. Several identified targets with major isoform switches represent established regulators of placental development, trophoblast migration, and vascular function, that are known to be altered in PE [14,160]. Such isoform switching could generate truncated or non-functional proteins or modify protein-protein binding affinities [161,162], representing a potentially under-researched pathological mechanism in PE that is well-documented in other disorders, such as tuberculosis or Alzheimer’s disease [163–165].

Finally, we evaluated differential DNAm patterns and regions (DMRs) in PE placentas, given that DNAm can repress gene expression by hindering transcription factor access or alter exon inclusion to determine the predominant gene isoforms [41]. However, we identified no significant hits, contrasting previous literature noting significant changes in DNAm in PE [110,166–168]. This contrast could reflect the heterogeneity of PE. Specifically, our compiled dataset lacked the metadata required to control for disease subtypes, likely masking DNAm shifts unique to these subtypes. Overall, our findings suggest that the involvement of DNAm in PE pathology and its downstream impact on gene expression is limited. The observed transcriptional changes are, therefore, likely driven by other forms of gene regulation, such as histone modifications, microRNAs, altered transcription initiation/termination sites, or post-transcriptional gene silencing and degradation [169].

### Shifts in placental cell-type proportions in PE

We performed cell-type deconvolution to determine whether shifts in placental cell-type proportions in PE could account for the observed transcriptomic. Female PE placentas exhibited higher proportions of fetal mesenchymal stem cells and fetal nRBCs relative to healthy controls. Although tissue processing aims to remove circulating blood, differences in detected nRBC levels between PE and controls may reflect true pathological elevations [121]. Elevated nRBCs may reflect a compensatory response to placental hypoperfusion and fetal hypoxia in PE, as these cells help maximize oxygen transport under ischemic stress [170,171]. Similarly, PE-associated hypoxia might stimulate MSC self-renewal to promote placental angiogenesis as a compensatory mechanism [172,173]. The DNAm-based deconvolution partially conflicted with the RNA-based findings, showing a lower abundance of stromal cells in PE placentas across both sexes. This difference could be due to cell-type resolution differences, with the "stromal cell" classification possibly encompassing fetal MSCs. However, the DNAm array did not detect variations in fetal nRBCs. These findings demonstrate that placental cellular compositions shift slightly in a pathology-dependent manner.

### Strengths, limitations, and future directions

The primary strength of our study lies in its multi-omics approach and high statistical power, analyzing combined public datasets across several genetic and epigenetic metrics. We also stratified analyses by sex, allowing us to isolate signals that would otherwise be masked in combined-sex cohorts. This study is also one of the first to perform alternative splicing and isoform switching analysis in PE, providing novel insights into transcriptional regulation as a driver of pathology. However, there were several technical limitations in our study. Compiling public datasets for a meta-analysis prevented us from controlling for confounding variables and pre-existing comorbidities, or uncovering differences between early- and late-onset PE, since metadata was not consistently available. In addition, because our DNAm and RNA expression populations were not from a matched single-cohort, true biological correlations may be underestimated. Lastly, since our analysis was performed on term placentas, our findings reflect late-stage PE pathology rather than its early development, preventing us from fully uncovering the mechanisms of lipid-related dysregulation.

Future work should more thoroughly characterize the relationship between alternate transcription sites and PE pathology to better understand potential disease mechanisms. Specifically, functional studies may help understand how isoform switches may drive pathology. In addition, future studies should aim to characterize the lncRNAs that consistently exhibited high differential expression in PE. Identifying their function may enhance our understanding of the mechanisms governing PE and reveal therapeutic targets. Finally, collecting first- or second-trimester cell-free DNA and RNA, will allow us to assess the utility of lipid-candidate genes as early biomarkers, and help map how dyslipidemia in PE evolves overtime.

## 5. Conclusion

Overall, our findings demonstrate that lipid dysregulation in PE placentas may stem from changes at the gene and transcript level, while DNAm shifts at these candidate loci are less prominent. Utilizing interaction term analysis, we identified several sex-specific DEGs in PE, although many were uncharacterized in the literature. We also observed a slight shift towards a male-specific downregulation of genes involved in aerobic metabolism, suggesting a preference of anaerobic metabolism in male-PE placentas. In addition, we identified several significant isoform switches that were entirely undetected by our differential gene expression analysis. Many of these switches occurred in pathways dysregulated in PE and were driven by alternative transcription initiation and termination sites rather than alternative splicing. A subset of these isoform switches yielded positive predicted functional consequences, highlighting how pathological mechanisms may be masked when analyzing gene-level abundance alone. Abnormal isoform expression may therefore contribute to PE pathogenesis, representing a novel research area. Ultimately, our findings suggest that PE-associated lipid and broader placental changes are driven primarily by transcriptional variations, and may serve as promising targets for future biomarkers and mechanistic research.

## 7. Data Availability

All source code can be found on the Wilson Pregnancy Lab Github [52]. All data are already on GEO, and how to extract those datasets is within the source code.

## 8. Author’s Roles

K.E., M.L., and S.L.W. all substantially contributed to the study conception and finalized manuscript. M.L. and X.Y. screen for publicly available data. K.E. completed all analyses and was responsible for drafting this manuscript. All authors contributed to manuscript revisions and approved the final version for submission. All authors agreed to be accountable for the accuracy and integrity of this work.

## Supplementary Figure Captions

Supplementary Figure 1: Flowchart outlining sample removal in RNA-seq pre-processing workflow.

Supplementary Figure 2: PCA plot of RNA-Seq samples from studies GSE143953, GSE148241, GSE186257, GSE234729 before (left) and after (right) controlling for batch-effects. GSE234729 formed two distinct, isolated clusters that could not be controlled for, resulting in its removal from the metadata sample set.

Supplementary Figure 3: PCA plot of RNA-Seq data before (left) and after (right) normalization with DESeq2. Plots show closer clustering of data points after normalization, indicating a reduction of batch effects.

Supplementary Figure 4: PCA plot of DNA methylation data before (left) and after (right) normalization with adjusted functional normalization. Plots show closer clustering of data points after normalization, indicating a reduction of batch effects.

Supplementary Figure 5: Volcano plot depicting X-Chromosomal differential lipid-related gene expression (RNA-Seq) between PE and control placentae in the (A) female-stratified population (n = 62; 22 PE, 40 Control), and (B) male-stratified population (n = 30; 17 PE, 13 Control).

Supplementary Figure 6: Volcano plot depicting X-Chromosomal differential genome-wide gene expression (RNA-Seq) between PE and control placentae in the (A) female-stratified population (n = 62; 22 PE, 40 Control), and (B) male-stratified population (n = 30; 17 PE, 13 Control).

Supplementary Figure 7: Volcano plot depicting X-chromosomal DNAm in lipid-candidate genes between PE and control placentae in the (A) female-stratified population (n = 83; 48 PE, 35 Control), and (B) male-stratified population (n = 89; 52 PE, 37 Control).

Supplementary Figure 8: Volcano plot depicting x-chromosomal DNAm genome-wide between PE and control placentae in the (A) female-stratified population (n = 83; 48 PE, 35 Control), and (B) male-stratified population (n = 89; 52 PE, 37 Control).

