## Supplementary figures and images for "“Transcriptional and isoform-level regulation of lipid-candidate genes in preeclamptic placentas”"

### Supplemental Figure 1

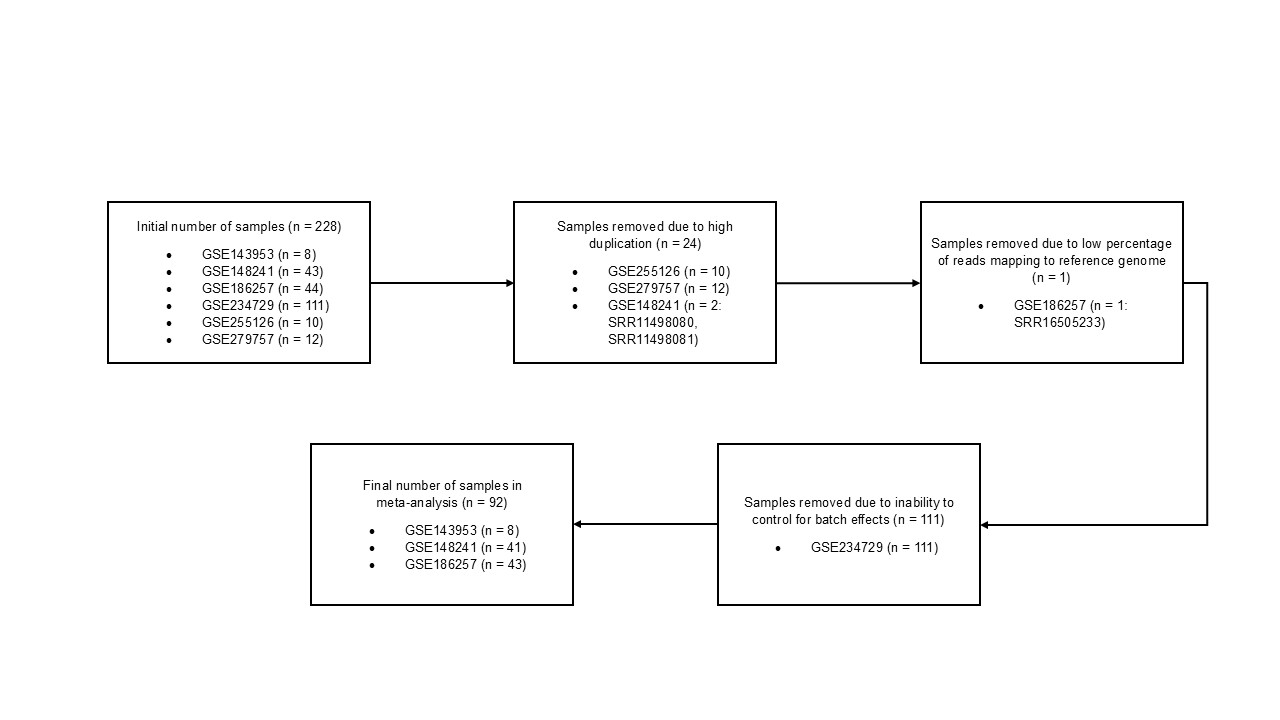

### Supplemental Figure 2

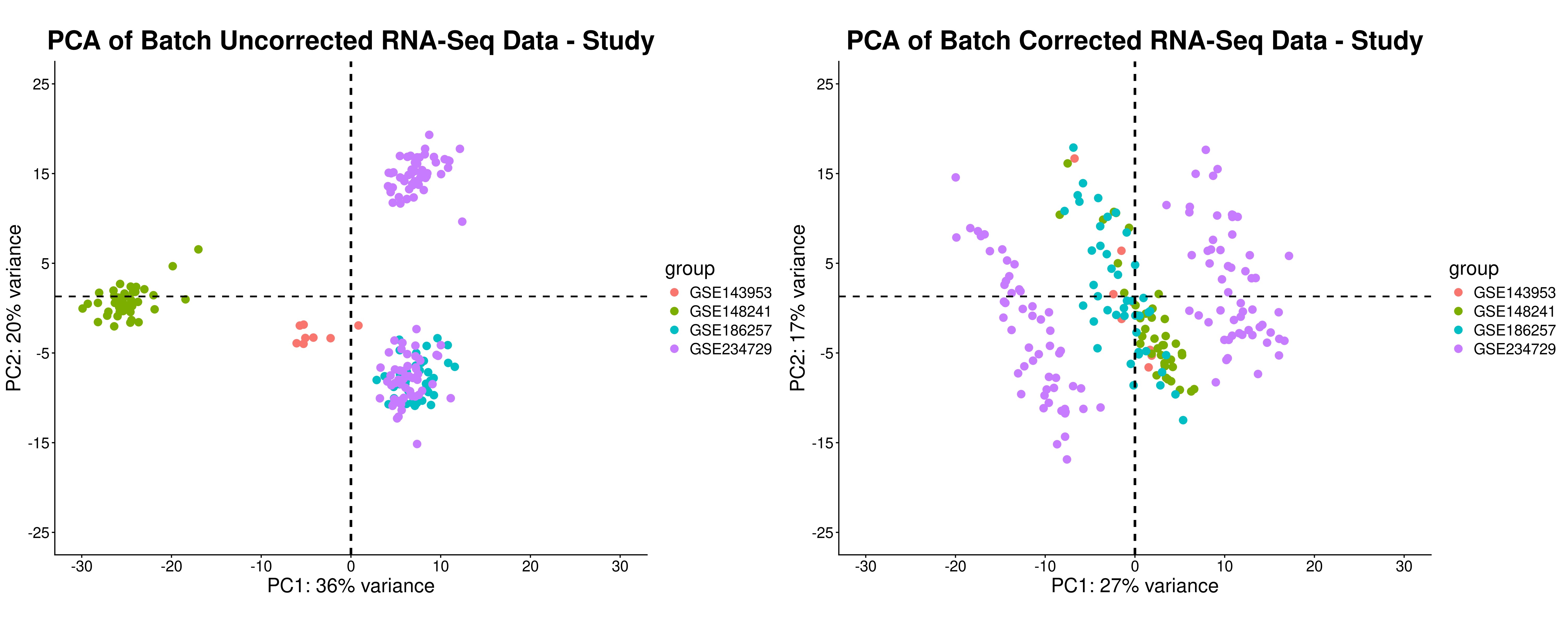

### Supplemental Figure 3

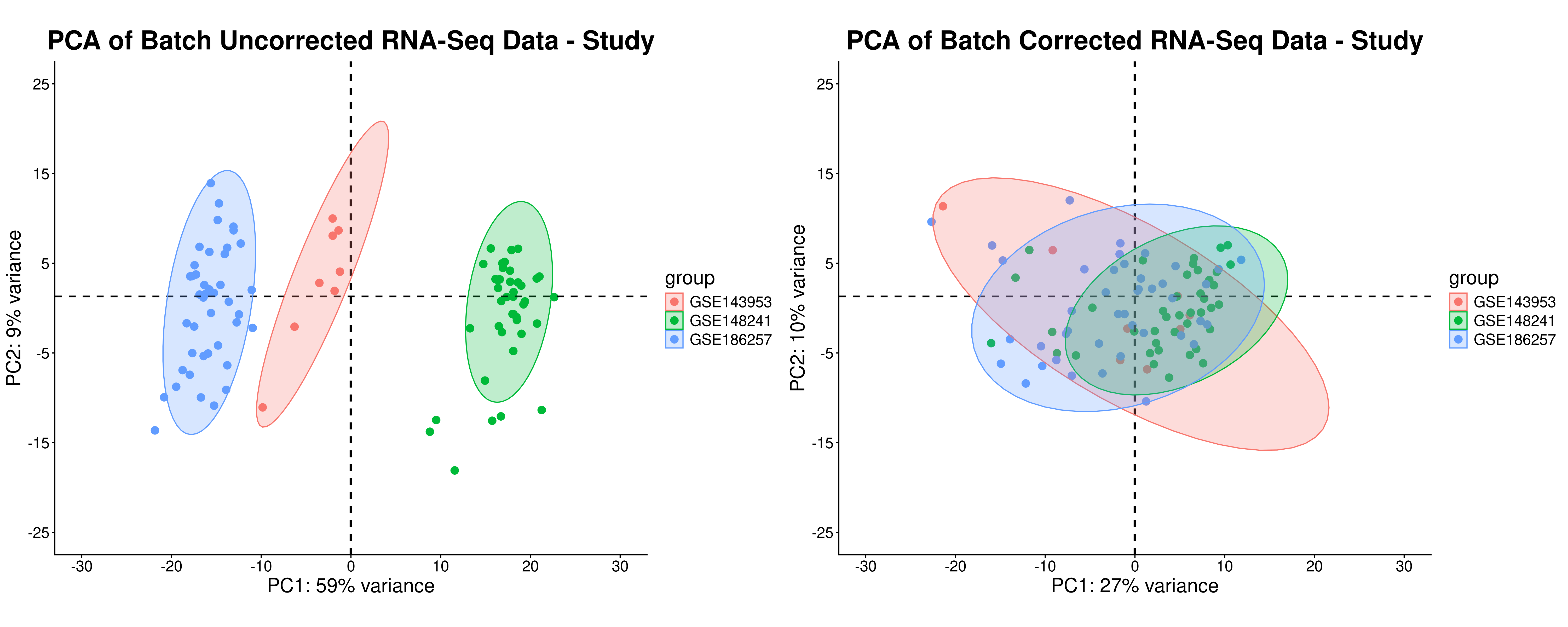

### Supplemental Figure 4

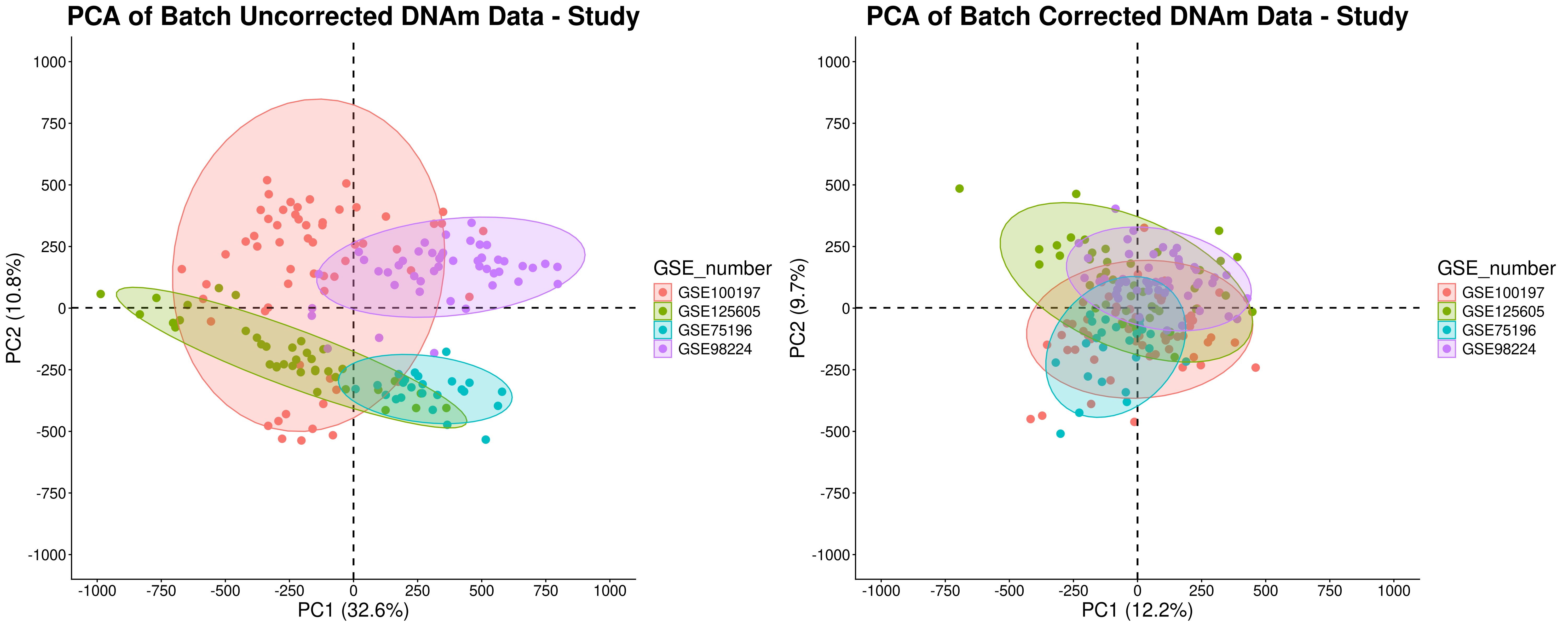

### Supplemental Figure 5

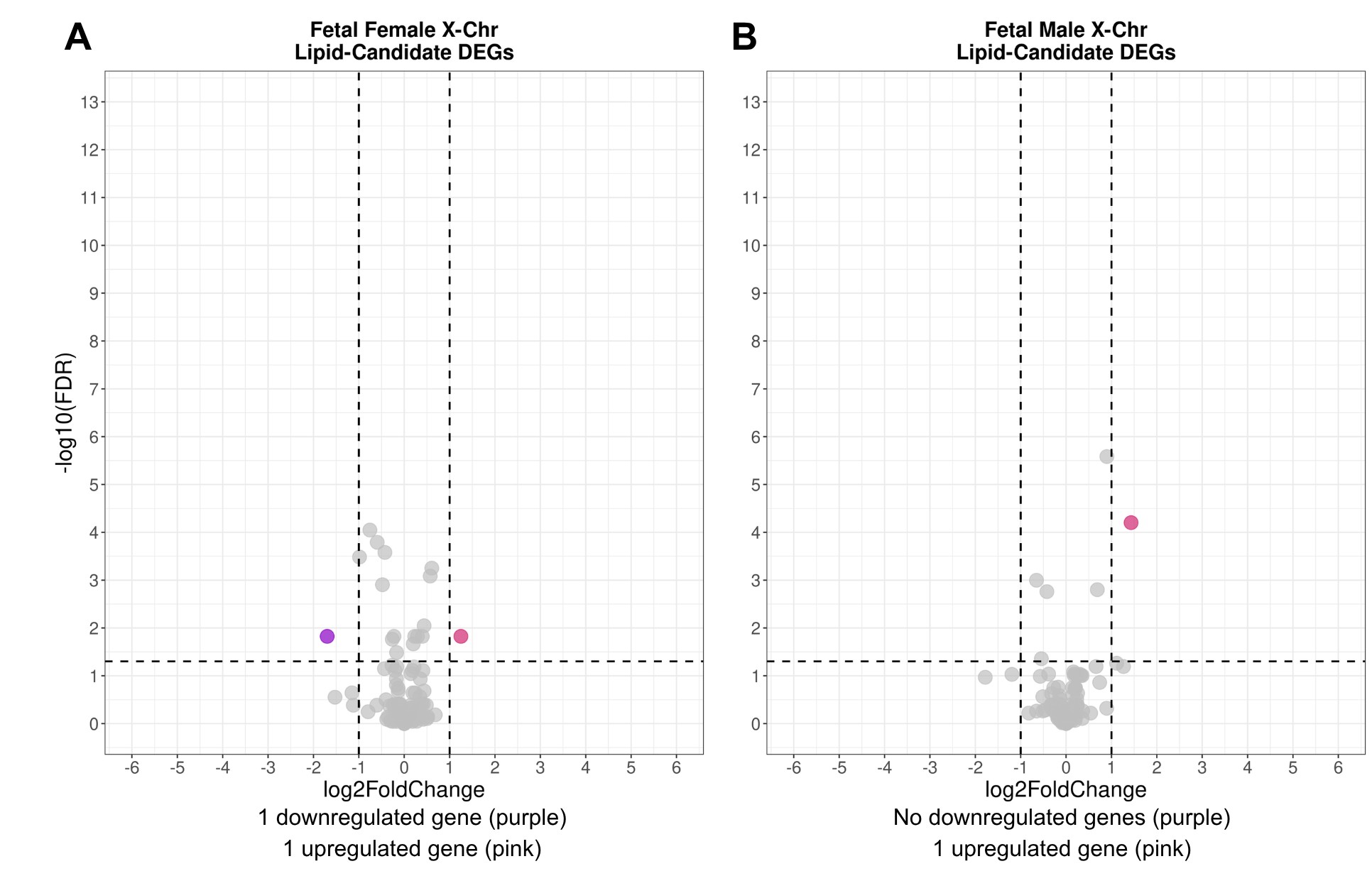

### Supplemental Figure 6

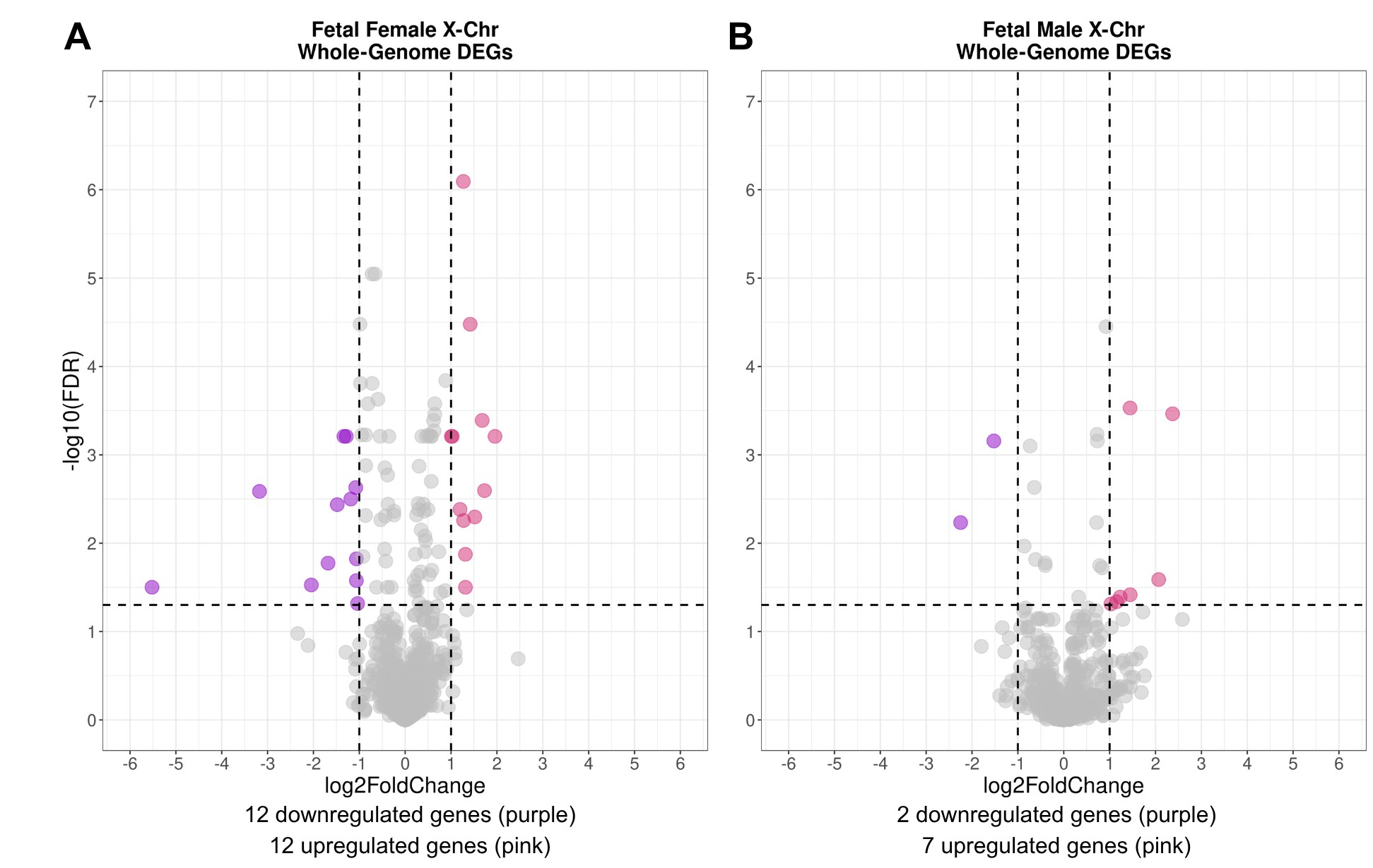

### Supplemental Figure 7

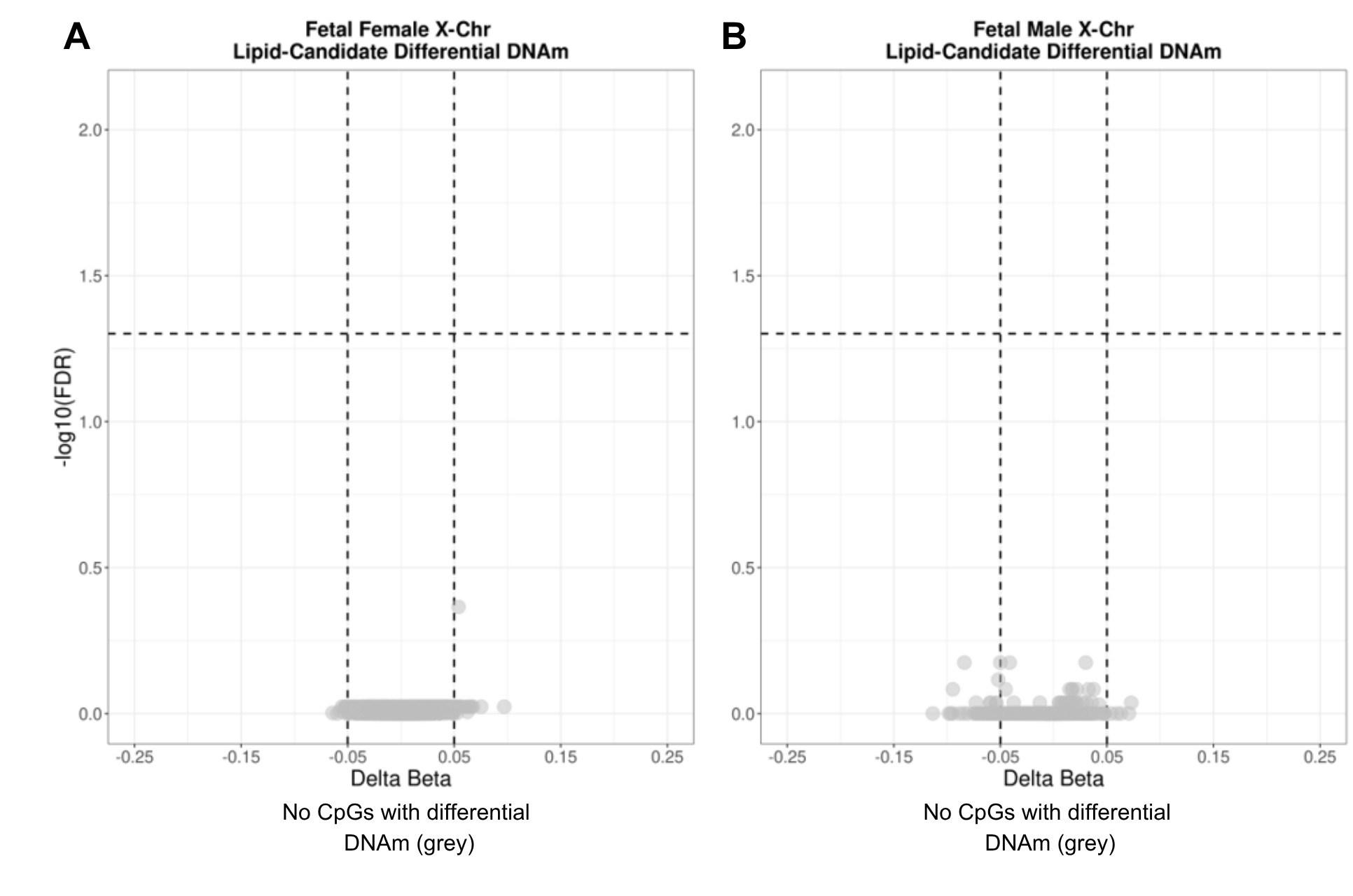

### Supplemental Figure 8

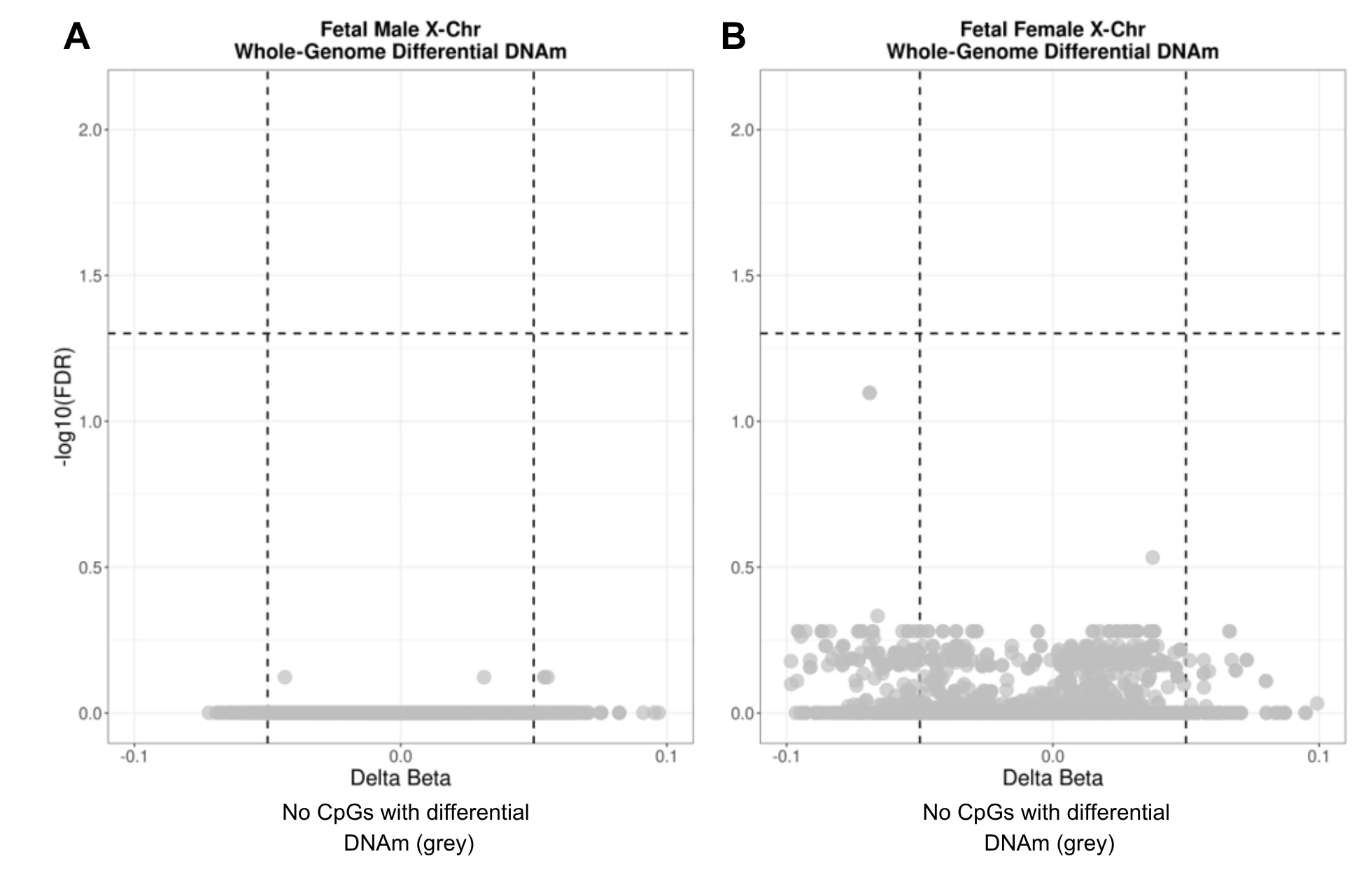
